# Controlling striatal function via anterior frontal cortex stimulation

**DOI:** 10.1101/191361

**Authors:** Mieke van Holstein, Monja I. Froböse, Jacinta O’Shea, Esther Aarts, Roshan Cools

**Affiliations:** Radboud University, Donders Institute for Brain, Cognition and Behavior, Nijmegen, The Netherlands; Department of Psychology and Brain Research Center, University of British Columbia, Vancouver, BC, Canada; Wellcome Centre for Integrative Neuroimaging (WIN), Oxford Centre for Functional MRI of the Brain (FMRIB), Nuffield Department of Clinical Neurosciences, University of Oxford, John Radcliffe Hospital, Headington, Oxford OX3 9DU, UK; Radboud University Medical Center, Department of Psychiatry, Nijmegen, The Netherlands

## Abstract

Motivational, cognitive and action goals are processed by distinct, topographically organized, corticostriatal circuits. We aimed to test whether processing in the striatum is under causal control by cortical regions in the human brain by investigating the effects of offline transcranial magnetic stimulation (TMS) over distinct frontal regions associated with motivational, cognitive and action goal processing. Using a three-session counterbalanced within-subject crossover design, continuous theta burst stimulation was applied over the anterior prefrontal cortex (aPFC), dorsolateral prefrontal cortex, or premotor cortex, immediately after which participants (N=27) performed a paradigm assessing reward anticipation (motivation), task (cognitive) switching, and response (action) switching. Using task-related functional magnetic resonance imaging (fMRI), we assessed the effects of stimulation on processing in distinct regions of the striatum. To account for non-specific effects, each session consisted of a baseline (no-TMS) and a stimulation (post-TMS) fMRI run. Stimulation of the aPFC tended to decrease reward-related processing in the caudate nucleus, while stimulation of the other sites was unsuccessful. A follow-up analysis revealed that aPFC stimulation also decreased processing in the putamen as a function of the interaction between all factors (reward, cognition and action), suggesting stimulation modulated the transfer of motivational information to cortico-striatal circuitry associated with action control.

## Introduction

The frontal cortex is responsible for many higher-order functions, such as goal setting, planning, and action selection. The frontal cortical regions involved in these functions are connected with anatomically distinct sub-regions of the striatum, organized in topographically specific circuits^1–4^. Connections between ventral/anterior parts of the prefrontal cortex (PFC) and the ventral/anterior caudate nucleus form a reward circuit, and have been associated with motivational goal setting^5^. Another circuit, connecting the dorsolateral prefrontal cortex (dlPFC) with more dorsal/posterior parts of the caudate nucleus, is associated with cognitive control processes^6^. Finally, in the action circuit, (pre-)motor cortices are connected with the posterior parts of the striatum, the putamen^7^.

Initial evidence for the existence of these topographically specific cortico-striatal circuits came from work with experimental animals^8–10^. Although this animal work generally converges with human work^3,11,12^, the majority of techniques (i.e. neuroimaging) used to investigate circuits in the human brain are correlational, and thus do not inform about whether activity in one region *causes* a change in another region. Several studies have used transcranial magnetic stimulation (TMS) to show that frontal regions can exert control over striatal processing. In the absence of any specific behavioral task, offline TMS applied over the dlPFC and the motor cortex affected processing in the striatum^13–16^. Such cortical control over striatal processing was shown to be functionally specific, at least in the motor domain: TMS over cortical motor areas changed processing in the putamen during tasks that depend critically on the motor circuit^17,18^. Whether motivational reward processing in the striatum is under control of (more ventral/anterior)cortical regions in the human brain remains unclear. Here we aimed to test this, while also assessing the functional and anatomical (topographic) specificity of such fronto-striatal control.

We employed an offline TMS protocol, continuous theta burst stimulation (cTBS)^19^, aimed at decreasing^15,20,21^ neural signaling in the three frontal regions embedded within functionally distinct cortico-striatal circuits known to implement motivational, cognitive and action goal processing. Using data from an independent fMRI dataset with the same paradigm^22^, we selected three cortical stimulation sites for this study. We did so by assessing the main effect of reward, switching between tasks (cognition), and switching between response buttons (action). Thus, a reward-related region in the aPFC was selected as being part of the reward circuit, a task-switch-related region in the dlPFC as part of the cognitive circuit, and a response-switch-related region in the premotor cortex (PMC) as part of the action circuit. TMS was followed immediately by task-related blood oxygenation level dependent (BOLD) functional magnetic resonance imaging (fMRI), to measure the consequences of these interventions on the task-evoked BOLD signal in the striatal subregions embedded within each functionally distinct circuit: the ventral/anterior caudate nucleus, dorsal/posterior caudate nucleus, and the putamen. Task-related processing was assessed using an established paradigm that we have used extensively to investigate reward anticipation (reward) and cognition (switching between tasks)^22–25^. To assess task-related processing in each of these cortico-striatal circuits, we stimulated these three cortical sites on three separate days, using a counterbalanced within-subject crossover design. Moreover, to account for non-specific effects related to the day ratherthan to the stimulation, each session consisted of two task-related fMRI runs: a no-TMS baseline and a post-TMS stimulation run (in counterbalanced order).

Based on *in vivo* evidence about the topography of human cortico-striatal connectivity^3,12^, we predicted that TMS over each cortical target site would attenuate functionally specific activity in its striatal target site. Specifically, we predicted a) that TMS over the aPFC would attenuate activity in the ventral/anterior caudate nucleus, the main striatal target of the aPFC specifically related to one functional domain (reward-processing), but not another (task switching or response switching). We furthermore set out to test the prediction that b) TMS over the dlPFC would attenuate activity related specifically to task switching in the posterior caudate nucleus, the main striatal target of the dlPFC^3,12^ and c) that TMS over the PMC would attenuate activity related specifically to switching between response buttons in the putamen, the main striatal target of the PMC^3,12^. In addition, we anticipated these effects to be anatomically specific at the level of the striatum: i.e., we anticipated that cortical stimulation of one frontal region (e.g. TMS over the aPFC) would alter processing in a distinct region of the striatum (e.g. caudate nucleus) and not in another region (e.g. putamen). Finally, we anticipated this effect would be anatomically specific at the level of the cortex: i.e., specific to TMS over one frontal region (e.g. TMS over the aPFC, but not of the dlPFC or PMC, would alter processing in the caudate nucleus).

We performed additional exploratory analyses of the effect of TMS over the aPFC. These analyses were based on the idea that the organization of cortico-striatal circuits is not strictly parallel. Instead, signals from each cortico-striatal circuit can be transferred to other circuits^3,8–10,26^, thereby providing a mechanism by which reward-predictive signals (in the reward circuit) can engage and alter cognitive processes (in the cognitive circuit) and guide action selection (in the action circuit). We tested this possibility in two ways. First, we assessed the effect of aPFC stimulation as a function of the two-way interaction between reward anticipation and task switching, expecting modulation in more dorsal/posterior parts of the caudate nucleus, which is part of the cognitive control circuit^6,22^. Second, we assessed the effect of aPFC stimulation as a function of the three-way interaction between reward, task switching and response switching, expecting to observe modulation in the sub-region of the striatum that is part of the action circuit, namely the putamen^3,7,18^.

## Materials and methods

### Participants

Of the 31 participants who started the main experiment, 27 participants (ranging from 18 – 25 (mean 21.7, SD 1.95) years old; 14 men) completed all sessions and were included in the analyses (**Supplementary Information**). Participants were TMS and MRI safety screened and gave written informed consent according to the Declaration of Helsinki and guidelines of the local ethics committee on research involving human participants (CMO Arnhem/Nijmegen: 2011/244). They received course credits and/or payment for their participation.

### Experimental design and procedures

All sessions took place at the Donders Centre for Cognitive Neuroimaging in Nijmegen, The Netherlands. The experiment consisted of four visits to the center: one ‘intake’ session and three experimental sessions.

#### Intake session

During the intake session, participants were introduced to the paradigm and completed two practice blocks on a laptop and a third practice block (**paradigm**) in the MRI scanner during the acquisition of a structural scan (**MRI procedure**). Next, in order to determine the stimulation intensity for each individual, we assessed their active motor threshold (aMT) (**TMS procedure**). Finally, participants were familiarized with the sensation of cTBS in order to ensure tolerability of cTBS over the stimulation sites (**TMS procedure**). After successful completion of the intake session, the first of three experimental sessions followed at least one week later.

#### Experimental sessions

During each experimental session, participants completed two runs in the MRI environment. To account for nonspecific effects, the paradigm was administered twice: once ∼10 minutes after TMS (stimulation fMRI) and once without the prior influence of TMS (baseline fMRI)^17^. Finally, to control for order effects, we counterbalanced the order in which the stimulation fMRI and baseline fMRI runs were completed (figure 1). Specifically, 14 participants first performed the baseline fMRI run, followed by TMS, followed by the second fMRI run (stimulation fMRI; figure 1a). The remaining 13 participants started with TMS and fMRI (stimulation fMRI), followed by a 30-minute break - to allow for the TMS effects to wear off - and then the second fMRI run (baseline fMRI; **figure 1b)** (**Supplementary Information**: **Order effects**).

**Figure 1.**
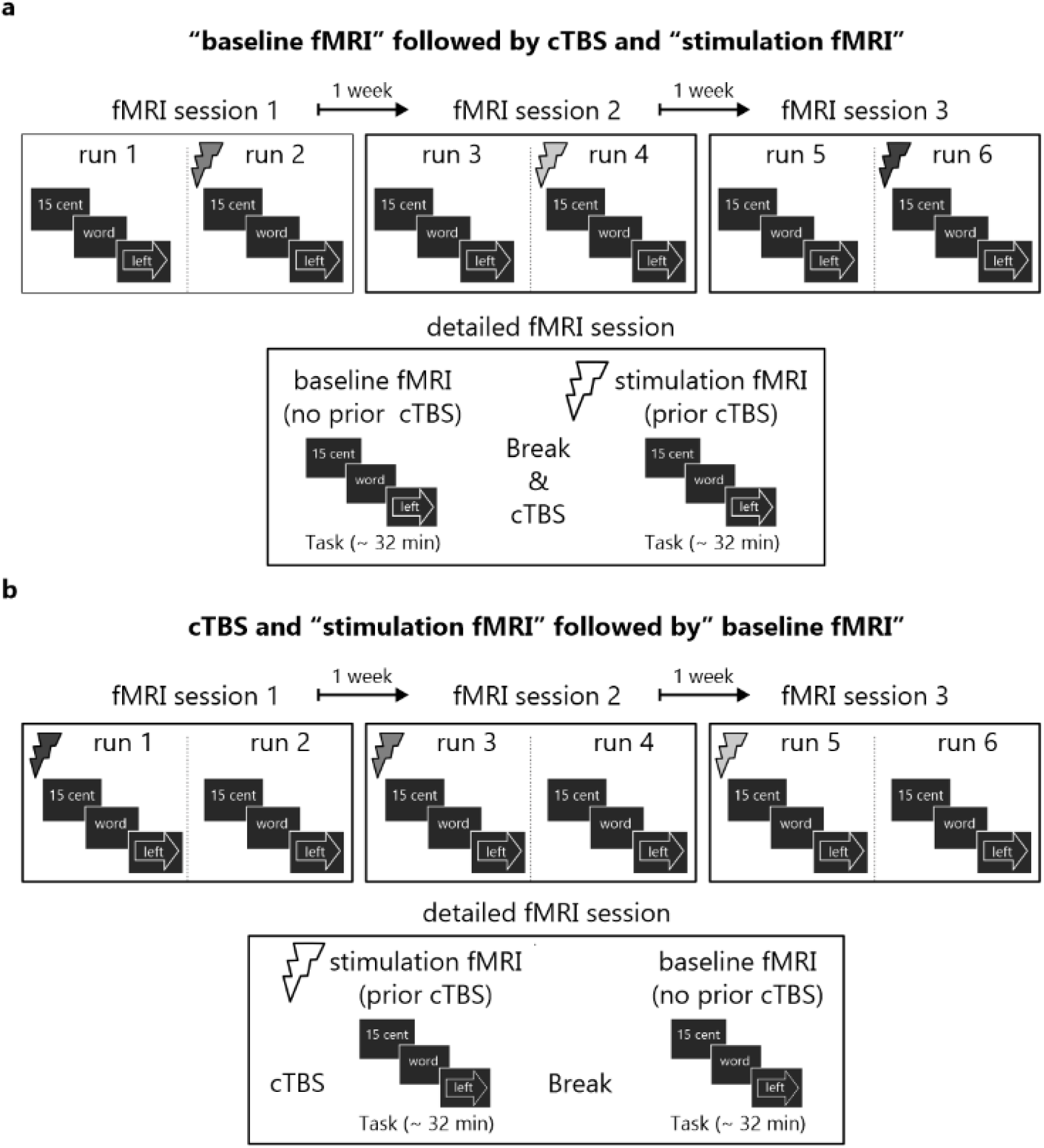
Experimental design. Each participant received continuous theta burst stimulation (cTBS) (indicated with thunderbolts) on three occasions. On each occasion, a different cortical region was targeted with TMS: anterior prefrontal cortex, dorsolateral prefrontal cortex, and dorsal premotor cortex (indicated with distinctly shaded thunderbolts in the **top panels of a & b**). The order in which each participant received cTBS over these regions was counterbalanced between participants. During each session, each participant completed two fMRI runs; one after TMS (stimulation fMRI) and one without prior influence of cTBS (baseline fMRI). The order in which a participant would perform the stimulation fMRI and baseline fMRI run was counterbalanced between participants (see **bottom panels of a & b** for a detailed illustration of an fMRI session). **a**) Halfthe participants (N=14) started with a baseline fMRI run, then received cTBS followed by the stimulation fMRI run. **b**) The remaining participants (N = 13) received the opposite arrangement; i.e. the session started with the application of cTBS and a stimulation fMRI run, followed by a baseline fMRI run. See table 1 for the times between cTBS and the fMRI scans.

Previous work has shown that cTBS over the motor cortex can suppress motor evoked potential (MEP) amplitudes for up to 50 minutes after stimulation, with effects no longer evident after 60 minutes^19,21^. Therefore, in the current study, for participants who did the stimulation fMRI run first, the delay between the administration of cTBS and the start of the baseline fMRI run was approximately 90 minutes (range: 87 – 107 minutes) (table 1).

**Table 1.** Time between onset of the two fMRI runs and between cTBS onset and fMRI run onset, for each session separately.

|  | aPFC | dIPFC | PMC | Average |
| --- | --- | --- | --- | --- |
| **Order 1 = Baseline fMRI – cTBS – stimulation fMRI** |  |  |  |  |
| cTBS – stimulation fMRI * | 8.73 ± 0.43 | 9.11 ± 1.10 | 9.21 ± 1.00 | 9.02 ± 0.70 |
| Baseline fMRI – stimulation fMRI * | 94.14 ± 5.60 | 92.86 ± 4.70 | 92.07 ± 4.84 | 93.02 ± 3.44 |
| **Order 2 = cTBS – stimulation fMRI – baseline fMRI** |  |  |  |  |
| cTBS – stimulation fMRI * | 10.02 ± 2.01 | 9.37 ± 0.90 | 9.47 ± 0.98 | 9.62 ± 1.06 |
| Stimulation fMRI – baseline fMRI * | 85.31 ± 11.16 | 83.08 ± 2.93 | 82.46 ± 3.26 | 83.62 ± 4.81 |
| cTBS – baseline fMRI * | 92.41 ± 5.48 | 92.94 ± 3.22 | 91.45 ± 3.13 | 92.26 ± 3.44 |

\*Differences are calculated from the start of the first fMRI run (duration: ~35 minutes) or the start of 40s cTBS, to the start of the second fMRI run (duration: ~35 minutes). Times displayed are average ± SD minutes.

### Paradigm

Participants performed a task-switching paradigm with a reward manipulation that has been used previously ^e.g. 25^. In the current version, minor changes were made to include a response-switching component. All details of the task are described in the legend of figure 2.

**Figure 2.**
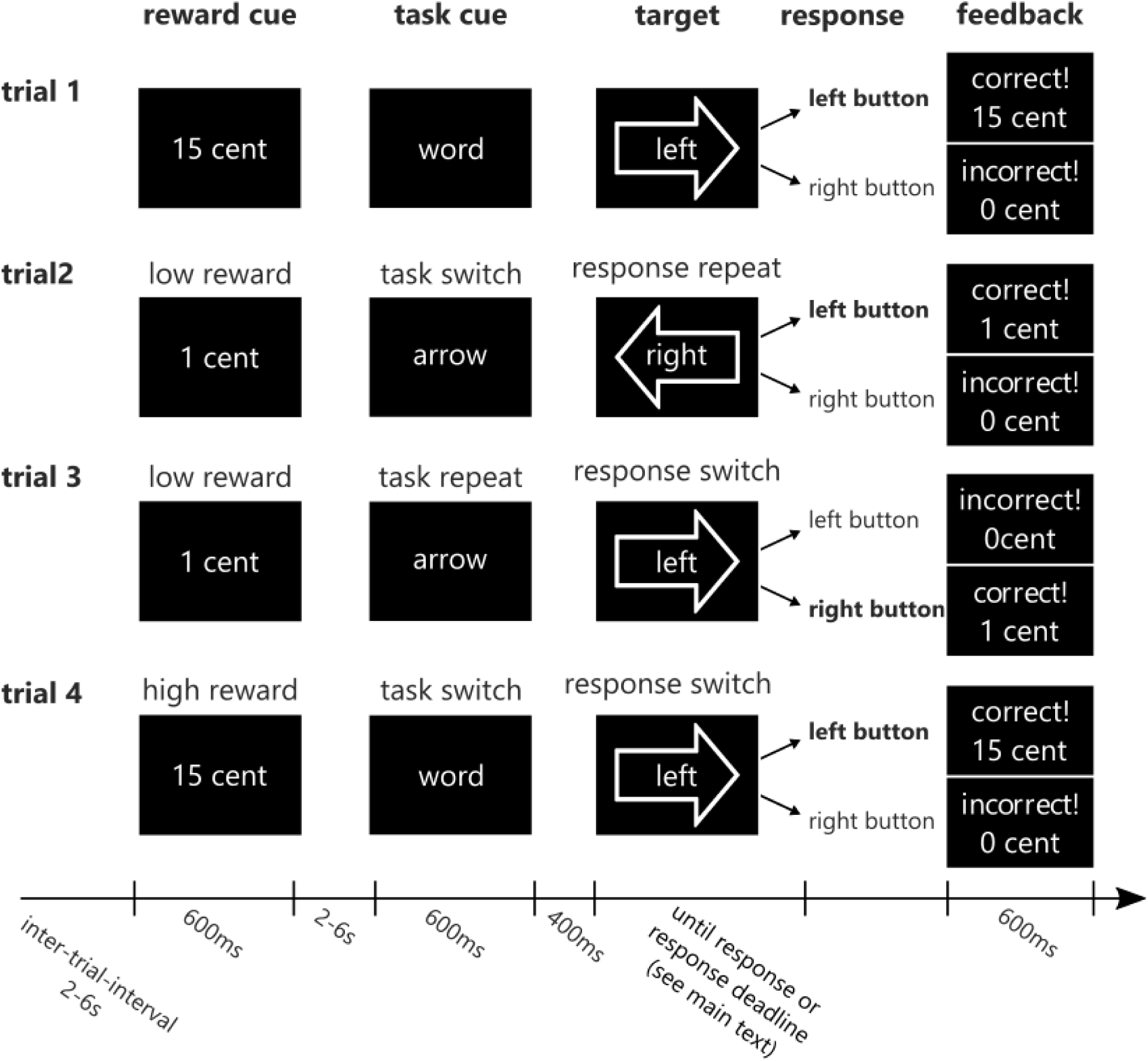
Task and response switching paradigm with reward manipulation. Participants always had to respond to response-incongruent arrow-word combinations (targets) with a left or right button press, either by responding to the direction indicated by the arrow (i.e. [←] or [→]) or to the direction indicated by the word (i.e. ‘left’ or ‘right’). A task cue preceding the target (by 400ms) indicated which task (arrow or word) the participant had to respond to on the current trial. The task on the current trial could either change (unpredictably) with respect to the preceding trial (i.e. task switch trial; arrow-word as in trial 4, or word-arrow as in trial 2) or remain the same (i.e. task repeat trial; arrow-arrow (trial 3), or word-word). In addition to such task switches, the paradigm allowed us to look at response switches, i.e. whether the correct response (left or right button, the correct response is printed in bold), remained the same (i.e. response repeat trial; left-left as in trial 2, or right-right) or switched (i.e. response switch trial; right-left, as in trial 4 or left-right as in trial 3) compared with the previous trial. In the current version of the paradigm we made sure that the task switches occurred independently from response switches; half of the task-repeat trials and half of the task-switch trials required a switch of the responsebutton (e.g. trial 3 and 4 respectively), whereas the remaining half of the trials required a response repetition (e.g. trial 2). In addition, as in previous versions, we manipulated the amount of anticipated reward (€0.01 vs. €0.15) on a trial-by-trial basis by means of a reward anticipation cue. At the start of each trial this reward cue indicated the amount of reward for that trial, contingent on a correct and sufficiently fast button press (see **paradigm**). Immediately following the response, feedback was given (e.g., “correct! 15 cents” in green ink, or “incorrect! 0 cents” in red ink) ^see also 25^.

Each session started with a number of practice blocks (**Supplementary Information**). The paradigm consisted of 160 trials and lasted ∼35 minutes with a 30s break every 32 trials. In the breaks and at the end of each run (i.e. after 160 trials) the cumulative amount of money the participant earned was displayed on the screen (max. €12.80). Participants were informed in advance that we would keep track of the total amount of money on each run and that the total earnings of one of the six runs would be added to their financial compensation as a bonus. Which run this was, was determined by chance at the end of the third session.

### Transcranial Magnetic Stimulation (TMS) procedure

#### Selection and targeting of stimulation sites

The mean MNI coordinates for targeting TMS at each of the three stimulation sites were determined by assessing the peak BOLD-fMRI activations for the main effect of Reward (high > low reward cue), the main effect of Task switching (task switch > task repeat), and the main effect of Response switching (response switch > response repeat) in the frontal cortex. This was performed on a previously published fMRI dataset that had used the same behavioral paradigm^22^. A region in the left anterior PFC (aPFC: −30, 60, 8, Brodmann area 10, **white circle in** figure 3a) was identified as part of the reward network; a region in the left dorsolateral PFC (dlPFC: −36,36, 20, Brodmann area 46, **white circle in** figure 3b) was identified as part of the cognitive (task switching) network; and a region in the left premotor cortex (PMC: −28, 10, 66, Brodmann area 6, **white circle in** figure 3c) was identified as part of the action (response switching) network.

**Figure 3.**
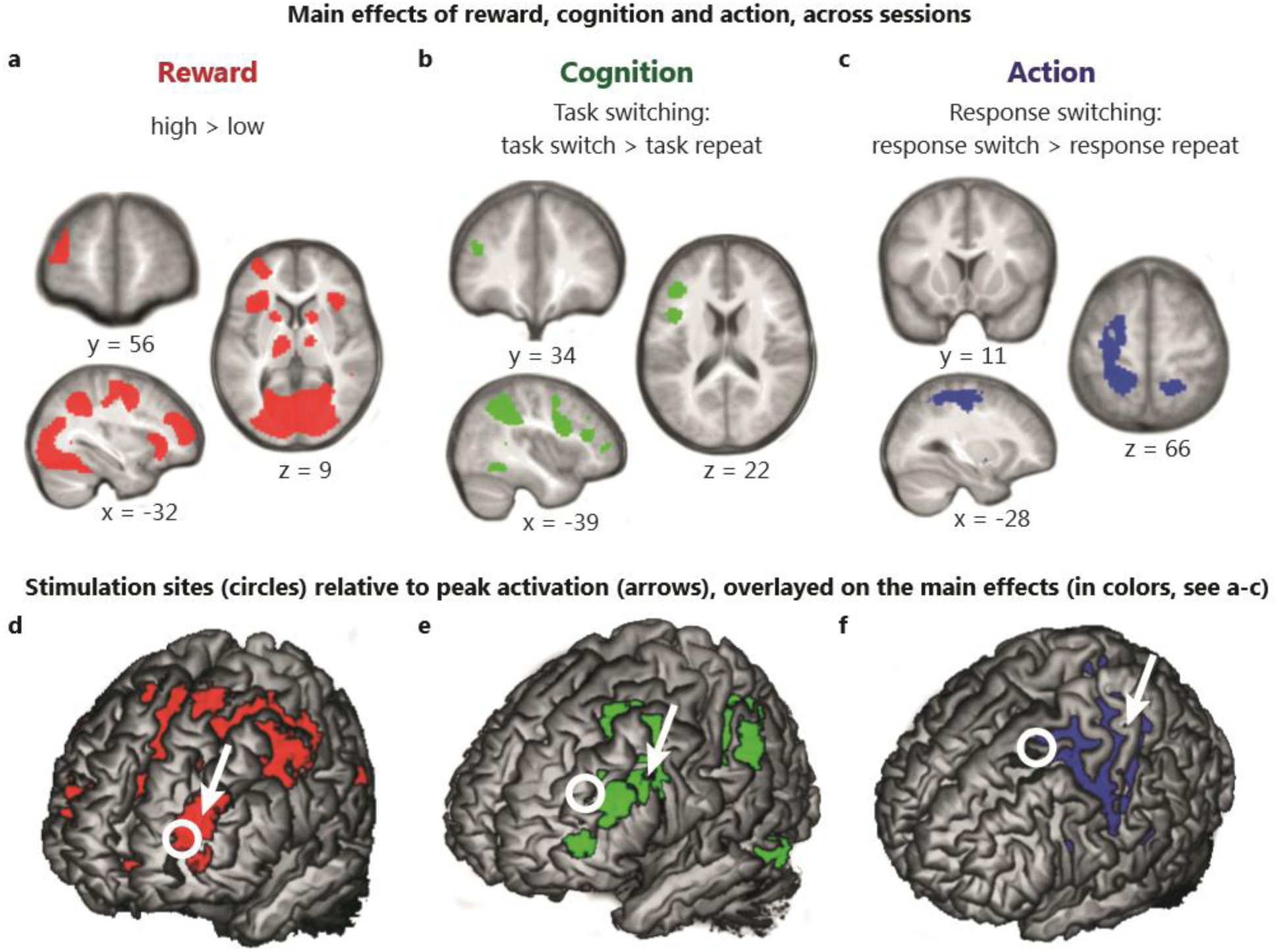
Main effects of task manipulations across runs. The three vertical panels show the main effect of: Reward anticipation (high reward > low reward; red; **a & d)**, task switching (task switch > task repeat; green; **b & e);** and response switching (response switch > response repeat; blue; **c & f)**, combined across all experimental runs. Results are displayed at a threshold of t = 3.14 (P_UNC_ < 0.001). See table 2 **and the main text** for whole-brain and FWE small-volume corrected results. For illustration purposes, white circles (8 mm diameter) indicate the coordinates at which stimulation was applied to the **d)** anterior prefrontal cortex, **e)** dorsolateral prefrontal cortex, and **f)** premotor cortex, determined based on an independent fMRI dataset^22^. Arrows point to the location of the peak activations nearest the stimulation site for the relevant factor of interest in the current study. The rendered images show regions with a search depth of 8 mm.

To determine the coil positioning, we converted the stimulation targets in group mean MNI coordinates into individuals’ native anatomical space (**Supplementary Information**).During the intake session, participants were familiarized with the sensation of cTBS over each of these regions by undergoing 10 seconds (instead of 40 seconds) of stimulation, using otherwise identical parameters to those used in the main experimental sessions (**Supplementary Information**).

**TABLE 2.** Main effect of Reward anticipation (across all sessions)

|  | Peak MNI coordinate |  |  | t-value | Statistic |  |
| --- | --- | --- | --- | --- | --- | --- |
|  | x | y | z |  | p-value (peak) | cluster size |
| Reward (high > low) |  |  |  |  |  |  |
| Frontal lobe |  |  |  |  |  |  |
| SFG/ MFG (B10) | -32 | 50 | 18 | 6.15 | P<sub>FWE</sub> < 0.001 | 977 |
| SFG: SMA (B6)<sup>BI</sup>, dACC (B32)<sup>BI</sup> | -8 | 4 | 60 | 8.79 | P<sub>FWE</sub> < 0.001 | 7318 |
| Subcortical |  |  |  |  |  |  |
| Striatum<sup>BI</sup>: caudate nucleus * | 10 | 10 | 0 | 8.11 | P<sub>FWE</sub> < 0.001 | 4543 |
| Thalamus | 24 | -24 | 4 | 4.92 | P<sub>FWE</sub> = 0.043 | 50 |
| Occipital lobe |  |  |  |  |  |  |
| Lingual gyrus (B17)<sup>BI</sup> | -12 | -94 | -4 | 16.18 | P<sub>FWE</sub> < 0.001 | 213500 |
| Parietal lobe |  |  |  |  |  |  |
| Posterior cingulate cortex<sup>BI</sup> (B23) | -4 | -30 | 26 | 5.47 | P<sub>FWE</sub> = 0.005 | 538 |
| Reward (low > high) |  |  |  |  |  |  |
| Frontal lobe |  |  |  |  |  |  |
| IFG (B10) | 46 | 40 | 2 | 5.22 | P<sub>FWE</sub> = 0.013 | 555 |
| MFG: OFC (B11) | -40 | 36 | -12 | 5.20 | P<sub>FWE</sub> = 0.015 | 403 |
| MFG (B9) | 32 | 32 | 48 | 5.07 | P<sub>FWE</sub> = 0.025 | 697 |

The table shows all areas that were significant at peak $P_{\text{FWE}} < 0.05$ .
\* Cluster includes the caudate nucleus, putamen, nucleus accumbens, midbrain, thalamus, pallidum and extends into the insular cortex and IFG; SMA = supplementary motor area, dACC = dorsal anterior cingulate cortex; OFC = orbitofrontal cortex; SFG: superior frontal gyrus; MFG: middle frontal gyrus; IFG: inferior frontal gyrus; BI = bilateral cluster; B = Brodmann area

#### Continuous Theta Burst Stimulation (cTBS) protocol

During the experimental sessions, cTBS was administered following the protocol described by Huang and colleagues^19^. They applied cTBS (bursts of three 50 Hz pulses every 200ms for 40s,i.e. a total of 600 pulse) over the motor cortex at 80% of the aMT and reported a depression of MEP amplitudes over a subsequent period up to 60 minutes ^see also 21^. We chose to use this stimulation protocol (cTBS) which inhibits motor corticospinal excitability, rather than TMS (e.g. 10Hz repetitive TMS, rTMS) - which is excitatory - for practical reasons. Although rTMS has been shown previously to modulate striatal processing, and so could have been used to address our research questions, a practical advantage of cTBS over rTMS guided our choice: When applied near facial muscles, cTBS over the aPFC is reportedly well tolerated^20^, contrary to anecdotal evidence regarding rTMS protocols.

TMS pulses (biphasic) were administered through a figure-eight coil (75mm diameter), connected to a MagPro X100 stimulator (Mag Venture, Denmark). We used standard electromyogram (EMG) recordings to visualize MEPs from the first dorsal interosseous muscle of the right hand and to determine the resting motor threshold, using a standard protocol^20,27^ (**Supplementary Information**).

### Magnetic Resonance Imaging (MRI) procedure

#### MRI acquisition

MRI images were acquired on a 3-Tesla MRI system (Magnetom TrioTim; Siemens Medical Systems, Erlangen, Germany), using a 32-channel head coil. High-resolution T1-weighted MP-RAGE anatomical images were acquired during the intake session (GRAPPA acceleration factor 2; repetition time 2300 ms; echo time 3.03ms; field of view: 256 mm; voxel size 1 mm^3^). In order to obtain a good signal-to-noise ratio for brain areas susceptible to dropout, functional images were acquired using a T2*-weighted multi-echo gradient-echo planar sequence (repetition time: 2090 ms; echo times for 4 echoes: 9.4, 21.2, 33, 45 ms; flip angle: 90°; 32 ascending slices; 0.5 mm slice gap; voxel size 3.5 × 3.5 × 3 mm)^28^. In addition, following each task-related fMRI acquisition, we acquired 266 resting state scans (data not reported).

#### Preprocessing of task-related fMRI data

All data were analyzed using SPM 8 (Statistical Parametric Mapping; Wellcome Department London, UK, http://www.fil.ion.ucl.ac.uk/spm). Prior to standard preprocessing, the four echo images were combined using echo summation (**Supplementary Information**). The combined images were slice-time corrected to the middle slice, normalized to a standard template, and smoothed (**Supplementary Information**).

### Statistical analyses

#### Statistical analysis of behavioral data

Behavioral analyses were performed on the response times (RTs) and error rates (**Supplementary Information**). Results were analyzed using a repeated measures ANOVA with the factors TMS condition (stimulation or baseline), Reward (high vs. low), Task switching (task switch vs. task repeat) and Response switching (response switch vs. response repeat). We report effect sizes for all significant effects (p < 0.05) using partial eta squared (η_p_^2^).

#### Statistical analysis of fMRI data

##### First-level analysis

The preprocessed fMRI time series were analyzed at the first level using one general linear model (GLM) for each participant, including all sessions. For each session, the following 26 task-related regressors were modeled at the onset of the stimulus (duration = 0) convolved with a canonical hemodynamic response function. The main effect of reward was modelled at the onset of the reward cue (high/low). For the main effects of task switching, response switching and the interaction between task factors, the following events were modelled at the presentation of the arrow-word target: [the type of reward trial (high/low) × the task cue (arrow/word) × task (switch/repeat) × response (switch/repeat)]. We additionally modelled the feedback (correct low/correct high/incorrect/too late), breaks (duration = 30s), the first trial of each block, andresponse omissions. In addition, we accounted for residual head motion, movement-related intensity changes and low-frequency signals (**Supplementary Information**).

To assess the main effects (Reward, Task switching, Response switching), we generated, for each participant, a contrast image at the first level. For the main effect of Reward (high > low reward cue), the effect was time-locked to the reward cue. The contrast images for the main effects of Task switching (task switch > task repeat) and Response switching (response switch > response repeat) were all time-locked to the presentation of the arrow-word target.

##### Second-level analysis

At the second level, the contrast images of each effect were subjected to a full factorial GLM, including all sessions of each participant. First, we assessed the main effect of each component of the paradigm (i.e. Reward, Task switching and Response switching) across all six sessions (i.e. irrespective of TMS) (figure 3). Secondly, we addressed our primary question: whether stimulation of the prefrontal cortex *influences* task-related striatal processing. To this end, we assessed whether stimulation of the aPFC, relative to baseline (i.e. the contrast aPFC^BASE-STIM^), changed Reward-related processing in the striatum. More specifically, we anticipated that stimulation of the aPFC (vs. baseline) would reduce the BOLD response exclusively as a function of the main effect of Reward, and not as a function of the main effect of Task switching or Response switching (functional specificity), selectively in the caudate nucleus and not in the putamen (anatomical specificity at the level of the striatum) and exclusively after stimulation of the aPFC and not after stimulation of the dlPFC or PMC (anatomical specificity at the level of the cortex) (see **Supplementary Information** for a more detailed description of these analyses). In addition, we assessed whether stimulation of the dlPFC (dlPFC^BASE-STIM^) and the PMC (PMC^BASE-STIM^) changed processing related to Task-switching and Response-switching in the striatum.More specifically, we predicted TMS would modulate the BOLD response selectively in a more dorsal/posterior portion of the caudate nucleus and in the putamen, respectively. Finally, we performed two post-hoc exploratory analyses to further explore the effects of aPFC stimulation. Here, we aimed to test whether stimulation of the aPFC would not only affect processing as a function of Reward, but also processing as a function of the integration between Reward and Task switching, or between Reward, Task switching and Response switching. This was based on the idea that the organization of cortico-striatal circuits is not strictly parallel, but that signals from each cortico-striatal circuit can be transferred to other circuits^3,8–10,26^. To address this, we generated contrast images at the first level for a 2-way interaction (between Reward and Task switching) and a 3-way interaction (between Reward, Task switching and Response switching), all time-locked to the presentation of the arrow-word target. Functional and anatomical specificity was assessed for any effects below P_FWE_ < 0.05 (**Supplementary Information).**

##### Statistical testing and data visualization

Effects that survived a family wise error (FWE) correction (P_FWE_ < 0.05) were considered significant. We assessed effects at the whole-brain level and for specific hypotheses regarding the caudate nucleus and putamen. Therefore, we applied a small volume correction (SVC) in the bilateral caudate nucleus and bilateral putamen to assess the effects of cTBS on each of the main effects (figure 4). To account for the two regions of interest (the caudate nucleus and the putamen), P_FWE_ effects below (0.05/2=) 0.025 were considered significant for the ROI analyses.

**Figure 4.**
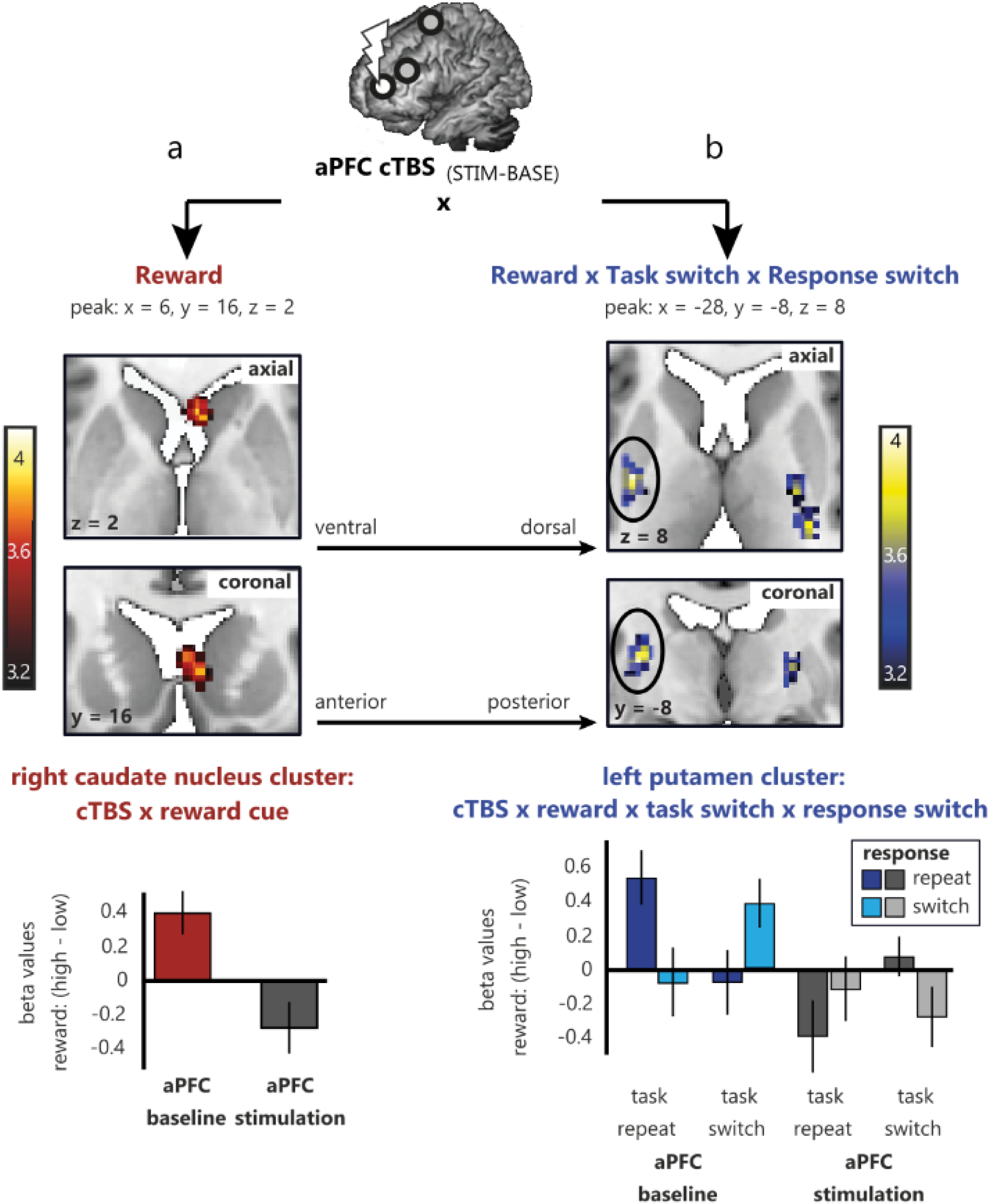
Effect of aPFC stimulation versus baseline for Reward (red) and for the interaction between Reward, Task switching and Response switching (blue) **top:** Brain maps for the effect of cTBS (_baseline - stimulation_) on the main effect of Reward (**a-** in red) and on the interaction between Reward, Task switching and Response switching (**b-** in blue) are shown at a threshold of PUNC < 0.001, t > 3.14. The FWE-corrected significant result (p<0.025 after Bonferroni correction for 2 ROIs) is indicated by the black circles. **Bottom**: Plots of the beta values extracted from the right caudate(**a**) and left putamen (**b**) cluster displayed in the top part of the figure. See **supplementary figure S1** for whole-brain maps across the axial (dorsal to ventral) and coronal (anterior to posterior) planeColor scales reflect t-values. Note that the results are displayed at a low threshold (PUNC < 0.001, t > 3.14), but that statistical significance (FWE-corrected) of the results was assessed in two anatomically defined regions of interest (i.e. restricted to the grey matter of the bilateral caudate nucleus and bilateral putamen). A high-resolution default MRIcron template was used as a background.

For visualization purposes, statistical maps are overlaid onto a study-specific template (**Supplementary Information**) and displayed at t = 3.14, p < 0.001, unless stated otherwise. Significant (FWE-corrected) results are reported in the text and tables and are indicated by black circles in figure 4. To visualize any effect that survived a P_FWE_<0.05 threshold, we extracted the beta values for the main effect of Reward and for the interaction between Reward, Task switching and Response switching from each activated cluster using the MarsBar function in SPM (figure 4, **Supplementary Information**).

## Results

### Effects of task on brain activity independent of cTBS

First, we assessed the main effect of Reward _(HIGH VS. LOW)_, Task switching _(SWITCH VS. REPEAT)_, and Response switching _(SWITCH VS. REPEAT)_, independent of cTBS, by pooling the data across runs (i.e. across three stimulation fMRI and three baseline fMRI runs).

Comparing the neural signal during high versus low Reward cues revealed a large bilateral network of regions, including the striatum, lingual gyrus, thalamus, cingulate cortex and the aPFC (table 2 and figure 3a). The opposite contrast (low – high) revealed three clusters in the prefrontal cortex (table 2)

During trials on which the Task switched, compared with trials on which the Task was repeated (Task _SWITCH-REPEAT_), a network of three clusters, encompassing the inferior frontal gyrus, cingulate gyrus, superior parietal lobe, and inferior temporal gyrus was activated (figure 3b).More specifically, the peak of one cluster was located in the inferior frontal gyrus (P_peak_FWE_ < 0.001, cluster size = 1755, T = 6.01, peak x, y, z = −40, 4, 30), one in the parietal lobe (P_peak_FWE_ < 0.001, cluster size = 3760, T = 8.72, peak x, y, z = −24, −66, 50) and one in the temporal lobe (P_peak_FWE_ < 0.001, cluster size = 751, T = 6.48, peak x, y, z = −48, −52, −12). A region of interest analysis with focus on the signal in the striatum did not reveal any significant effects of Task switching in either the caudate nucleus or the putamen. No regions exhibited increased activation during the opposite contrast (Task _REPEAT - SWITCH_).

One large cluster was more active during Response switching compared with Response repetition trials (Response _SWITCH-REPEAT_; figure 3c). This left lateralized cluster included the primary motor cortex (B4), premotor cortex (B6), primary somatosensory cortex, precentral gyrus and the primary somatosensory cortex (B3) and extended posteriorly into the parietal lobe, i.e. the postcentral gyrus (P_peak_FWE_ < 0.001, t = 6.61, z = 6.19, cluster size = 3250, peak x, y, z = −40, − 36, 54). A region of interest analysis with focus on the signal in the striatum did not reveal any significant effects of Response switching in either the caudate nucleus or the putamen. No regions exhibited increased activation during the opposite contrast (Response _REPEAT - SWITCH_).

In summary, analyses of the fMRI activation patterns (independent of TMS) revealed a significant main effect of Reward anticipation _(HIGH vs. LOW REWARD)_ in the caudate nucleus. However, no significant voxels were detected in the striatum as a function of the main effect of Task switching _(SWITCH –REPEAT)_ or the main effect of Response switching _(SWITCH –REPEAT)_, i.e. across all sessions. Therefore -in supplementary analyses-we investigated the data from each of the three fMRI sessions separately, to determine whether there was a main effect of Task switching during the baseline run of the dlPFC session, and a main effect of Response switching during the baseline run of the PMC session. These analyses also did not reveal any significant clusters in the striatum. The absence of effects in the striatum undermines the logic of testing whether cTBS decreases the BOLD response in the striatum during the stimulation vs. baseline run if no significant activation is detected during the baseline run. In addition, activated voxels for the main effect of Task switching were located up to 3.2cm away from the dlPFC stimulation site, and the activated voxels for the main effect of Response switching were located up to 4.6cm from the PMC stimulation site. By contrast, the stimulation coordinate for the aPFC was located within 1cm of the cluster that was activated by the main effect of reward. Perhaps not surprisingly then, analysis of the effect of cTBS over the dlPFC (_BASE-STIM_) on Task switching (_SWITCH – REPEAT_) or of cTBS over the PMC (_BASE-STIM_) on Response switching (_SWITCH – REPEAT_) did not reveal any brain regions where cTBS changed the BOLD-fMRI signal. Therefore, we will focus on results obtained after stimulation of the aPFC.

In what follows, we assessed our key hypothesis for the one session where the task evoked a significant effect on the BOLD response in the striatum: the aPFC, and asked whether1) changes in task-related processing occurred in the striatum after stimulating the aPFC, and whether these effects were 2) anatomically and 3) functionally specific.

### Effects of anterior prefrontal cortical stimulation on striatal processing

#### Functionally-specific effects of aPFC stimulation on striatal processing

Reward-related BOLD signal in the right caudate nucleus was decreased after aPFC stimulation compared with baseline (aPFC_BASE-STIM_ × Reward _HIGH-LOW_: P_SVC_FWE_ = 0.040, k = 17, T = 3.78, z = 3.69, peak x, y, z = 6, 16, 2; figure 4 – **in red**). The effect was located in the anterior portion of the caudate nucleus, and there was no such effect of aPFC stimulation on reward-related signal in the putamen, or elsewhere in the brain. However, this effect did not survive the Bonferroni correction (p<0.025) that we applied to the small volume corrected results in order to account for two striatal search volumes (bilateral caudate nucleus and putamen). Although we do not consider these results significant after correcting for multiple comparisons, for completeness, we report the assessment of functional and anatomical specificity in the **Supplementary Information**.

#### Transfer of task-related signal across cortico-striatal circuits

In an additional analysis we tested whether information from the Reward circuit was transferred across cortico-striatal circuits. This idea was motivated by anatomical work which has challenged the idea that cortico-striatal circuits are strictly parallel ^3,8–10,26^. Instead, according to theseproposals, signals from each circuit can be transferred to other circuits. We reasoned that – if these circuits are not strictly parallel - stimulation of the aPFC could affect task-related activity in more posterior parts of the striatum during the integration of Reward and Task switching or during the integration of all three factors: Reward, Task switching and Response switching.

There was no effect of aPFC stimulation on the integration of Reward and Task switching (i.e.: no 3-way interaction between aPFC_BASE-STIM_ × Reward _HIGH vs. LOW_ × Task _SWITCH vs. REPEAT_ (i.e. irrespective of Response switching). However, aPFC stimulation did modulate the integration between all three factors (i.e.: the 4-way interaction of aPFC_BASE-STIM_ × Reward _HIGH vs. LOW_ × Task _SWITCH vs. REPEAT_ × Response _SWITCH vs. REPEAT_ was significant). Assessment of the effect of aPFC stimulation on this integration (4-way interaction: aPFC_BASE-STIM_x Reward _HIGH vs. LOW_ × Task _SWITCH vs. REPEAT_ × Response _SWITCH vs. REPEAT_) revealed that aPFC stimulation decreased signaling in the left putamen (figure 4b) (left: P_SVC_FWE_ = 0.020, t = 3.96, k = 66; z = 3.86, peak x, y, z = −28, −8, 8). This effect remained significant after correcting for the two search volumes (i.e., p < 0.025). The signal in the right putamen was not significant (P_SVC_UNCORR_ < 0.001, k = 1). To illustrate these effects, the whole-brain maps at an uncorrected threshold of p<0.001 are shown in **supplementary figure S1**. To investigate further the nature of this interaction, one-sample t-tests were conducted on the Reward × Task × Response contrast images separately for each fMRI run.

Analysis of the aPFC baseline fMRI run revealed a significant Reward × Task × Response effect in the left putamen (left putamen: P_SVC_FWE_ = 0.01, t = 4.95, k = 64, z = 4.12, peak x, y, z = − 30, −18, 6; right putamen: P_SVC_FWE_ = 0.086, t = 4.21, k = 13, z = 3.64, peak x, y, z = 28, −12, 10) (figure 4b - **blue bars**). This was driven by a Reward effect on trials where the Task and Response either both switched, or both repeated, i.e. Task-Response congruent trials, with no Reward effect on Task-Response incongruent trials. By contrast, this 3-way interaction effect (Reward × Task × Response) in the left putamen was not significant during the aPFC stimulation run (figure 4b - **grey bars**), consistent with an inhibitory effect of aPFC TMS on striatal activity. Stimulation of the aPFC did not result in any other significant changes in BOLD signal elsewhere in the brain as a function of the interaction between Reward, Task switching and Response switching (**supplemental figure S1**).

To determine the anatomical specificity of this TMS effect at the level of the striatum, a direct comparison between the beta-values in the anatomically defined left putamen and the left caudate nucleus during the integration between Reward, Task switching, and Response switching was performed. This revealed that the TMS (aPFC _BASE-STIM_) effect in the putamen was anatomically specific: as evidenced by a significant ROI (left putamen vs. left caudate nucleus) × Stimulation (aPFC_BASE-STIM_) × Reward × Task × Response interaction: F(1,26) = 6.533, p = 0.017, η_p_ ^2^= 0.201.

To determine the anatomical specificity of this TMS effect at the level of the cortex, we assessed whether the reduction in BOLD signal related to the 3-way interaction between Reward, Task and Response during the aPFC session was statistically different from the stimulation vs. baseline effect during the other two (i.e. dlPFC and PMC) sessions (**Supplementary Information**). This analysis revealed one significant cluster in the left putamen: P_SVC_FWE_ = 0.0248, t = 3.92, z = 3.83, peak x, y, z = −26, −8, 12) **(Supplementary figure S3**). This finding shows that the effect of aPFC stimulation on task-related processing in the striatum was not likely related to non-specific effects such as the sensation of cTBS.

In summary, stimulation of the aPFC modulated activity in the posterior portion of the putamen as a function of the interaction between Reward, Task switching and Response switching (P_SVC_ FWE_ = 0.020). This effect was anatomically as well as functionally specific, both at the level of the cortex and the striatum.

### Behavior

#### General task effects, irrespective of TMS

A detailed description of the behavioral effects across all three sessions can be found in the **Supplementary Information, including Supplementary figure S4.** In summary, across sessions, we observed a main effect of Reward, Task switching, and Response switching. In addition, we observed an effect of Reward on Task switching, and between Task switching and Response switching.

#### No effects of TMS on behavior

We did not observe any significant main effects of stimulation, irrespective of task conditions (in response times or error rates: F(1,26 < 1). None of the main effects (i.e. of Reward, Task or Response) (**Supplementary Information**) were modulated by cTBS (stimulation vs. baseline) for any of the stimulation sites: response times and error rates all F(1,26) < 3.612, all p > 0.05. Finally, there was no effect of stimulation on the Reward × Task × Response effect for any of the stimulation sites (in response times and error rates: all F(1,26) < 2.316, all p > 0.1).

## Discussion

Functionally homogenous regions of the PFC and striatum are anatomically organized in parallel cortico-striatal circuits, connecting – for example – the anterior PFC and the anterior caudate nucleus as part of a reward circuit, and the motor cortices with the putamen as part of an action circuit. Moreover, the reward, cognition and action circuits are organized in an anterior to posterior and ventral to dorsal gradient, both in the frontal cortex and the striatum^3,11,12^. In the current study we aimed to assess these cortico-striatal circuits by asking whether task-related processing in the striatum is under control of the functionally homogenous region of the prefrontal cortex. For example, we aimed to assess whether reward-related processing in the caudate nucleus is under control of processing in the anterior PFC. To assess these hypothesized cortico-striatal causal interactions, we employed an offline TMS protocol aimed at decreasing neural signaling. In a three-session baseline-controlled counterbalanced within-subject crossover design, cTBS was applied over three distinct cortical sites, associated with motivational reward processing (anterior PFC), cognitive control (dorsolateral PFC) and action control (premotor cortex). Immediately following the application of cTBS, participants performed a rewarded task-switching paradigm in the MRI scanner. The paradigm tapped into all three cognitive domains: reward processing, task switching (cognitive control) and response switching (action control), allowing the assessment of task-related processing in distinct sub regions of the striatum (the caudate nucleus and the putamen).

We first assessed whether stimulation of the aPFC (compared with baseline) altered reward-related processing in the caudate nucleus. To correct for the comparison between twosearch volumes (caudate nucleus and putamen), we considered an effect P_SVC-FWE_< 0.025 as significant. We observed a marginal decrease (P_SVC-FWE_ = 0.040) in the reward-related BOLD response in the striatum after aPFC stimulation compared with baseline. However, this effect did not reach significance according to our statistical threshold. Accordingly, we question its reproducibility, particularly given the relatively low physiological plausibility of the effect (with the cluster extending into the cerebrospinal fluid) and the lateralization to the right, non-stimulated hemisphere.

Next, we tested whether information from the reward circuit (i.e. the aPFC) could be transferred to another cortico-striatal circuit. We observed that stimulation of the aPFC altered processing in the putamen, the sub-region of the striatum that is part of the action circuit, but only when assessed as a function of the interaction between reward, cognitive control and action control. Specifically, aPFC stimulation elicited a reduction in task-related BOLD response (compared with the aPFC baseline fMRI run). Importantly, we also found that this aPFC _(BASE vs. STIM)_ effect was different from the effect in the dlPFC _(BASE vs. STIM)_ and PMC _(BASE vs. STIM)_ sessions. This is an important finding, because it shows that the effect of aPFC stimulation on task-related processing in the striatum was not likely related to non-specific effects such as the sensation of cTBS. These findings provide causal evidence in humans that stimulation of the aPFC causes a change in task-related processing in the striatum and, moreover, show that such distally induced task-related modulation is not restricted to the targeted cortico-striatal circuit (reward), but that modulated processing in the reward circuit can alter processing in another circuit (i.e. in the action circuit). Previous work has shown that cTBS over the aPFC can alter processing in regionsdistant from the site of stimulation^20^. However, in that study perfusion-MRI, instead of BOLD was used. In addition, the effects of aPFC stimulation were observed in the amygdala during the processing of emotional stimuli. We show here that the more commonly used BOLD-fMRI technique can be used successfully to assess cTBS-induced changes in task-related striatal processing.

The current study does not directly address the mechanisms by which aPFC stimulation affects processing in the putamen. However, based on previous work, two potential routes are most plausible. First, signals from the aPFC could reach the putamen via direct cortico-striatal connections^10–12,26^. Second, the signal from the aPFC could be conveyed to the putamen via spiraling dopaminergic connections between the striatum and the midbrain, i.e. striatal-nigro-striatal (SNS) connections^8,9,29–31^. Although results are largely consistent across species, it should be noted that these theories are derived primarily from work with non-human primates or rodents.

The anatomical specificity of the effects at the level of the cortex should be interpreted with caution. It cannot be argued that specifically aPFC stimulation, and not stimulation of the other two regions, can affect the integration between reward, cognition and action in the putamen. For this claim to hold, it would be necessary to show that stimulation of the dlPFC or PMC was effective. The absence of any task-related effects on striatal processing after stimulation of the dlPFC or PMC in this study thus precludes this claim. The null-effect following stimulation of the dlPFC and PMC was unexpected, especially considering previous worshowing that stimulation of the dlPFC and the motor cortex can alter processing in thestriatum^15,17,18^. However, contrary to those other studies, the current study is the first to assess the effect of cTBS on task-related processing in the striatum. Although Ko and colleagues^15^ applied cTBS over the prefrontal cortex, they did not assess task-related processing, nor did they assess effects with BOLD-fMRI. While two other studies did observe effects of cortical TMS on task-related BOLD-fMRI in the striatum^17,18^, those studies did not stimulate the aPFC, but the primary motor cortex, using a different TMS protocol (rTMS: 1-10 Hz instead of cTBS).

One explanation of the null effect after dlPFC and PMC stimulation is that - irrespective of TMS - we did not observe an effect of task switching or response switching in the striatum. This was true both across all sessions, and during the relevant baseline session (e.g. the baseline dlPFC session for the effect of task switching). The logic of testing whether stimulation of a cortical region will reduce striatal BOLD response does not hold unless there is a measurable BOLD response in the striatum during baseline. Alternatively, it is conceivable that the cortical regions we stimulated do not have a (causal) role in switching between tasks (dlPFC) or response buttons (PMC). This is consistent with the distance between the peak of the task-related fMRI activity (**white circles in** figure 3d-f) and the stimulation sites (**arrow in** figure 3d-f). Stimulation was targeted based on fMRI coordinates from a previous study^22^. However, the coordinates functionally activated in the present study, both for task-switching (dlPFC) and response switching (PMC), differed from the previous sample to the extent that no task-related activation actually overlapped with the coordinates at which TMS was applied. Although the exact spatial resolution of TMS is highly dependent on several factors, a resolution in the range of millimeters up to several centimeters has been suggested^32^. In any case, the field induced by TMS declines rapidly with distance from the coil^33^. In the current study, this implies that we might have failed to stimulate the task-relevant circuits for the dlPFC and PMC but not for the aPFC sessions. More specifically, a meta-analysis of fMRI coordinates associated with task switching has identified the inferior frontal junction as a key node^34^. The region reported in the meta-analysis (x, y, z coordinates: −40, 4, 30) overlaps with a cluster activated by the task-switching contrast in the current study (x, y, z coordinates: −40, 4, 30). This same cluster in the inferior frontal gyrus has been observed in two previous studies using the same paradigm (x, y, z coordinates: −52, 6-, 34^24^; x, y, z coordinates: −48, 12, 28^25^). In combination, this suggests that the main effect of task switching in the inferior frontal gyrus observed in the current study was most likely a reliable effect, and that the dlPFC region we stimulated was too anterior to target the cortico-striatal circuitry crucial for task switching.

In terms of response switching, previous fMRI studies assessing effects of response switching embedded in a task-switching paradigm^35,36^, reported effects more posterior (y = 0) than our stimulation coordinate (y = 10). Interestingly, both studies reported peak coordinates that are located within the parietal cluster we report, suggesting that this may have been a more successful target for cTBS in the current study.

We previously reported modulation of the BOLD response in dorsal parts of the striatum during the integration between reward and task switching^22^. This interaction was not observed in the current dataset. An important feature of the previous work^22^ however was that those effects were revealed exclusively as a function of inter-individual variance in the dopamine transporter (*DAT1*) genotype^22,25^.

The current study was designed to modulate processing in the striatum. In keeping with this hypothesis, we showed that stimulation of the aPFC modulates BOLD responses in the striatum. However it did not induce any behavioral changes, which is not uncommon with offline TMS^17,37 (but see15,18)^. The absence of a behavioral effect precludes us from making any claims about whether stimulation of the aPFC and the consequent effects on the striatum were behaviorally relevant. On the other hand, it simplifies the interpretation of the observed BOLD changes. If stimulation had also changed behavior, this in itself would be expected to alter task-related signal in the striatum, complicating interpretation. In the absence of such effects, the observed changes in brain activity can be straightforwardly attributed to a causal effect of cortical TMS.

Several neuropsychiatric/neurological disorders, such as substance use disorder, obsessive-compulsive disorder, schizophrenia and Parkinson’s disease are accompanied by altered processing in cortico-striatal circuits ^for reviews see 38,39^. In addition, patients with Parkinson’s disease, obsessive-compulsive disorder, and depressive disorder can benefit from deep brain stimulation of subcortical regions such as the striatum^40–42^. Alternatively, TMS provides a potentially less-invasive method to target subcortical structures. In fact, TMS is currently being used as an FDA approved treatment for major depressive disorder and has been suggested as a potential treatment for patients suffering from schizophrenia^43^. The current work adds to the knowledge required for a better understanding of ways by which subcortical circuits can be targeted effectively via non-invasive cortical stimulation. We show that stimulation with cTBS, which is well tolerated over the anterior prefrontal cortex, can alter processing in the striatum.

## Acknowledgements

The authors would like to thank Ivan Toni and Inge Volman for advice regarding the design of the study. The authors would also like to thank Lennart Verhagen for technical TMS support and advice regarding data analysis. Finally, the authors would like to thank Paul Gaalman for technical MRI support.

## Author Contributions

JO’S, EA and RC designed the study. MvH and MF recruited volunteers and acquired the data. MvH analyzed the data. MvH, MF, EA, JO’S and RC wrote the manuscript.

## Additional Information

The authors declare that they have no competing financial interests. This work was supported by a Cognitive Neuroscience TOPtalent PhD scholarship to MvH, by the Netherlands Organization for Scientific Research (MaGW open competition grant by NWO awarded to RC, EA, and JO’S, 404-10-062), a Royal Society Dorothy Hodgkin Fellowship to JO’S, and a James McDonnell scholar award to RC (220020328).

## Corresponding author

Correspondence to Mieke van Holstein

## Supplementary Information

### Supplementary methods

#### Participants

One participant was excluded due to a contra-indication for MRI, one participant’s session was discontinued due to dizziness during MRI, one participant was excluded due to technical TMS problems, and one participant due to technical MRI problems. All 27 included participants had normal or corrected-to-normal vision, were right-handed and pre-screened for claustrophobia, psychiatric, neurological, and vascular disorders, drug and medication use, alcohol consumption and smoking behavior, as well as any contraindications for TMS and MRI.

### Paradigm

#### Practice blocks

The first practice block (24 trials), which was only administered during the intake session, included only task-switching to familiarize participants with the alternation between tasks (arrow vs. word). During this block, the task (i.e. whether to respond to the arrow or the word) alternated unpredictably from trial to trial without any reward cues, and the feedback on each trial was either “correct” or “incorrect”. During the intake session and at the start of each experimental session, participants completed a second practice block that was exactly the same as the actual paradigm described in the legend of figure 2, only shorter (i.e. 24 trials). Finally, a third block (32 trials) without reward or feedback was administered in the scanner immediately before the actual paradigm started, and during the intake session (see main text: **intake session**). This third block was used to determine each individual’s response window. We calculated the average response times on four trial types (arrow, word × task-switch, task-repeat), during the third practice block. These response times were set as the response deadline during the subsequent run. This was done to account for inter-individual and inter-run differences in response speed and subsequent task difficulty.

### Transcranial Magnetic Stimulation (TMS) procedure

#### Selection and targeting of stimulation sites

To determine the TMS coil positioning for each individual and each brain region, each participant’s structural scan was coregistered to the standard SPM8 T1 template (Montréal Neurological Institute; MNI) and segmented using a unified segmentation procedure^27^. This procedure resulted in a set of inverse parameters allowing the conversion of the stimulation targets in group mean MNI coordinates into individuals’ native anatomical space. Next, the MNI coordinates for each cortical stimulation site were projected onto each individual’s structural scan using a frameless stereotactic neuronavigation system (Localite, Sankt Augustin, Germany). The TMS coil was then placed on the scalp overlying the target coordinates (aPFC, dlPFC, PMC) using the Localite software.

During the intake session, participants were familiarized with the sensation of cTBS over each of these regions. Thirty-nine participants started the intake session. Any participant who reported - or showed - signs of discomfort during this part of the intake session was excluded from further participation. This resulted in the exclusion of eight participants: six participants due to discomfort during stimulation over the aPFC and two due to a more general feeling of discomfort during this part of the intake. As a result, 31 participants started the main experiment (see **participants**).

#### Continuous Theta Burst Stimulation (cTBS) protocol

During the determination of the active motor threshold (aMT), participants rested their right hand on a pillow while squeezing a small roll of tape with a pincer grip at 20% of their maximum strength, contracting their first dorsal interosseous FDI muscle continuously. The aMT was defined as the lowest stimulation intensity over the contralateral motor cortex that elicited reproducible MEPs (in at least 5 out of 10 successive stimulations). The aMT was 24%-37% (mean 30.44%, SD 3.61) of the maximum stimulator output.

### Magnetic Resonance Imaging (MRI) procedure

#### Preprocessing of task-related fMRI data

Prior to standard preprocessing, realignment was performed using the estimated head motion parameters (least-squares approach, 6 parameters) for the images with the shortest echo, which were applied to echo images for each excitation. The images of all sessions were aligned to the shortest echo of each session, and to the first session. After spatial realignment, the four echo images were combined using echo summation. The combined images were slice-time corrected to the middle slice and segmented using a unified segmentation procedure^27^. The bias corrected T1 image was coregistered to the mean functional image and the transformation matrix from the segmentation procedure was used for normalization to a standard template (MNI). Normalized images were smoothed using an 8 mm full-width half maximum kernel. A study– specific T1 template was generated from an average of all co-registered and normalized T1 images to display the results, using MRIcron software.

### Statistical analysis of fMRI data

#### First-level analysis

To account for residual head motion, the six original head motion parameters (3 translation, 3 rotation), their first derivative and the square of the original and first derivative were included in the model, resulting in 24 motion nuisance regressors^30^. In addition, we used the mean signal from the white matter and CSF to account for movement-related intensity changes^31^. Finally, a high-pass filter (128s) was used to remove low-frequency signals (e.g. scanner drifts) and an AR(1) model was applied to adjust for serial correlations in the data. Microtime onsets were adjusted to account for the earlier mentioned slice time correction.

**Supplementary figure S1.**
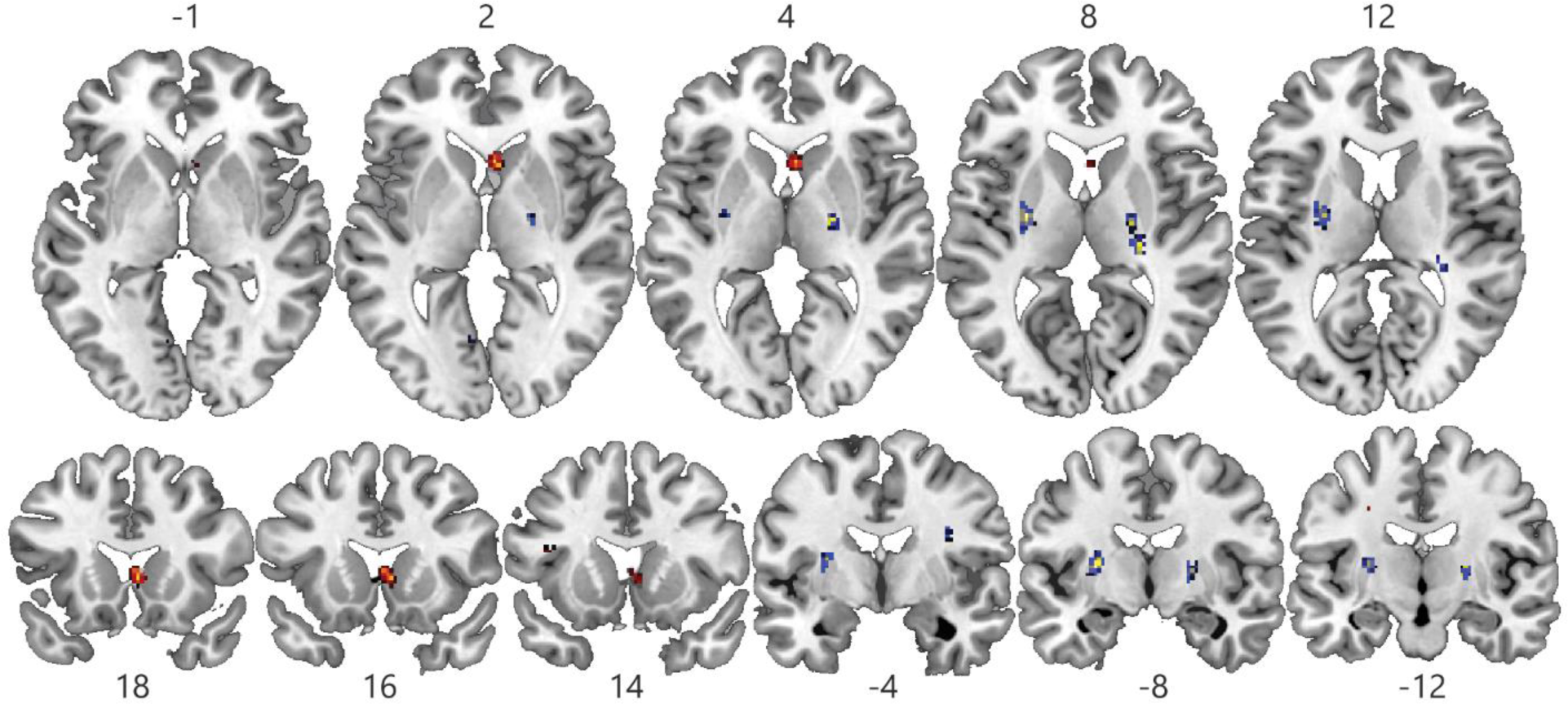
Whole-brain maps for the effect of aPFC stimulation on the main effect of Reward (in red) and the interaction between Reward, Task switching and Response switching (blue). Whole-brain maps across the axial (dorsal to ventral) and coronal (anterior to posterior) plane are shown for the effects in figure 4. The effect of aPFC stimulation on the main effect of Reward is shown in red. The effect of aPFC stimulation on the interaction between Reward, Task switching and Response switching in blue. Note that the results are displayed at a low threshold (PUNC < 0.001, t > 3.14), but that statistical significance (FWE-corrected) of the results was assessed in two anatomically defined regions of interest (i.e. restricted to the grey matter of the bilateral caudate nucleus and bilateral putamen).

### Functional and anatomical specificity: Supplementary methods and results

#### Functional specificity

To assess functional specificity, we tested whether TMS over the aPFC altered processing in the caudate nucleus exclusively as a function of Reward processing. In other words, we anticipated that TMS over the aPFC would not alter processing in the caudate nucleus as a function of Task switching or Response switching.

In addition, we assessed whether TMS over the aPFC altered processing in the putamen exclusively as a function of the interaction between Reward, Task switching and Response switching, and not as a function of any other effects (e.g. the effect of Reward, Task switching, *or* Response switching, or the interaction between Reward and Task switching).

The effects of aPFC stimulation on reward-related processing in the caudate nucleus and the effects of aPFC stimulation on the interaction between Reward, Task switching and Response switching in the putamen were functionally specific: aPFC stimulation did not decrease striatal processing as a function of Task switching (caudate nucleus: k = 0, putamen: P_SVC_-FWE_ = 0.165, k= 2) or Response switching (caudate nucleus: P_SVC_-FWE_ = 0.150 k = 3, putamen: k = 0). In addition, the effects of aPFC stimulation did not decrease striatal processing as a function of the interaction between Reward and Task switching.

#### Anatomical specificity at the level of the striatum

We assessed quantitatively the anatomical specificity of an effect in a region. For example, we aimed to assess whether the effect of aPFC stimulation on the effect in the left putamen (figure 4b) was different from the same effect in the left caudate nucleus. To avoid double dipping ^32^, we derived beta values from an independent anatomical ROI (i.e. independent from the activated cluster). Values were entered into a repeated measures GLM in SPSS with the factors stimulation (aPFC_STIM vs. BASE_), ROI (caudate nucleus vs. putamen) and either 1) Reward _(HIGH VS. LOW)_(time-locked to the reward cue) or 2) Reward _(HIGH VS. LOW)_, Task _(SWITCH VS. REPEAT)_ and Response _(SWITCH VS. REPEAT)_ (all time-locked to the target).

A direct comparison between the effect of Reward in the right caudate nucleus and the effect of reward in the right putamen revealed that the (trending) effect in the right caudate nucleus was specific to the caudate nucleus: ROI (right caudate nucleus vs. right putamen) × Stimulation (aPFC_BASE-STIM_) × Reward: F(1,26) = 4.937, p = 0.035, η_p_^2^=0.159

The effect in the putamen during the integration of Reward, Task and Response was anatomically specific: these results are discussed in the main text.

#### Anatomical specificity at the level of the cortex

To test whether the effect of aPFC stimulation on Reward-processing in the caudate nucleus was anatomically specific at the level of the cortex, we submitted the data from each session for the factor Reward to a GLM with the additional factor Site (aPFC, dlPFC, PMC).

This allowed us to assess whether the (marginal) effect of aPFC stimulation _(BASE-STIM)_ on reward-related processing in the caudate nucleus (figure 4a) was different during the aPFC session compared to the other two session. This interaction test (Stimulation × Site (i.e. aPFC _BASE-STIM_ > dlPFC _BASE - STIM_ = PMC _BASE − STIM_) × Reward) revealed no significant voxels in the whole-brain or the after applying a SVC in the striatum.

To test whether the effect of aPFC stimulation on the interaction between Reward, Task and Response was significantly different compared to the data from the dlPFC (_BASE − STIM_) and PMC (_BASE − STIM_) sessions, we submitted the data from each session for the Reward × Task × Response contrast to a GLM with the additional factor Site (aPFC, dlPFC, PMC), resulting in the following interaction test: Stimulation × Site (i.e. aPFC _BASE − STIM_ > dlPFC _BASE − STIM_ = PMC _BASE − STIM_) × Reward × Task × Response. This analysis revealed one significant cluster in the left putamen: P_SVC-FWE_ = 0.0248, t = 3.92, z = 3.83, peak x, y, z = −26, −8, 12). The whole-brain map for thisinteraction is shown in **figure S3a** at a threshold of P_UNCORRECTED_ < 0.001. To visualize this interaction we extracted the beta values as described elsewhere (see **order effects**). For an unbiased representation of the data, we extracted the beta values from the left anatomically defined putamen. We plotted the interaction between Reward, Task switching and Response switching on the Y-axis of **figure S3b**, separately for each stimulation site (aPFC, dlPFC, PMC) and for the stimulation fMRI vs. baseline fMRI run.

#### Order effects: Methods

We aimed to assess whether the results in the caudate nucleus (main effect of reward) and putamen (the Reward × Task switching × Response switching interaction), were dependent on the order in which participants performed the baseline and stimulation fMRI run (Stimulation order), i.e. stimulation fMRI followed by a baseline fMRI (N=13) or the opposite arrangement (N=14) (figure 1). We reasoned that any residual effect of the inhibitory TMS protocol in those who performed the stimulation fMRI run first, would be evident in a reduced effect during the baseline fMRI run in this group of participants. In addition, we aimed to assess whether we could find any evidence to suggest that the order of the stimulation Site (Site order), i.e. on which day the aPFC stimulation took place (i.e. during the 1^st^, 2^nd^ or 3^rd^ session), had an effect on the Reward effect or on the interaction between Reward, Task switching, and Response switching.

We extracted the beta values during the Reward cue (high and low Reward) from the cluster in the right caudate nucleus as presented in figure 4a. Next, using SPPS software (IBM SPSS Statistics 23), we performed a repeated measures GLM with the within-subject factors Stimulation (aPFC stimulation vs. aPFC baseline), and Reward (high vs. low) and the between subject factors Stimulation order and Site order (as described above).

From the cluster in the left putamen, as presented in figure 4a, we extracted the beta-values during target for the interaction between Reward, Task switching and Response switching and entered these variables as well as Stimulation (aPFC stimulation vs. aPFC baseline) as within-subject factors. We added the between subject factors Stimulation order and Site order (as described above).

In addition, because we observed a Site (aPFC, dlPFC, PMC) × Stimulation (stimulation vs. baseline) × Reward × Task × Response interaction in the putamen (**figure S3a**), we also assessed whether that interaction varied either as a function of Stimulation order or Site order. Because the voxels from which the beta values were extracted were based on the interaction between aPFC _BASE-STIM_, we assessed these effects in an unbiased region: the anatomically defined left putamen.

### Order effects: Results

#### The effect of aPFC stimulation on Reward processing in the caudate nucleus

We did not find any evidence to suggest that the effect of aPFC stimulation vs. aPFC baseline on the (trending) Reward-related signal in the caudate nucleus (figure 4a) was different in those participants who started the session with the baseline fMRI run (**figure S2a** dark red bars: baseline followed by stimulation) compared with those who started the session with cTBS followed by the stimulation fMRI run (**figure S2a** light red bars: simulation followed by baseline). More specifically, the Reward × Stimulation _(_aPFC _BASE-STIM)_ × Baseline order interaction was not significant (F(1,21)<1. In addition, there was no evidence that the effect of aPFC stimulation vs. aPFC baseline on the Reward-related signal in the caudate nucleus was dependent on the session number (1^st^, 2^nd^ or 3^rd^) in which the aPFC was stimulated (F(2,21) = 2.351, p >0.05.

#### The effect of aPFC stimulation on the integration between Reward, Task and Response in the putamen

Inspection of **figure S2b** suggests that the effect of aPFC stimulation vs. baseline on the integration between Reward, Task switching and Response switching (figure 4 – in blue) was smaller in participants that started the session with the baseline fMRI run (**figure S2b** darker blue bars), compared with the other group (**figure S2b** lighter blue bars). However, this effect was not significant: The effect of aPFC stimulation vs. aPFC baseline on the integration between Reward, Task, and Response in the putamen was not different for those participants who started the session with the baseline fMRI run (**figure S2b** dark blue bars: baseline followed by stimulation) compared with those who started the session with cTBS followed by the stimulation fMRI run (**figure S2b** light blue bars: stimulation followed by baseline). More specifically, the Stimulation (aPFC _BASE-STIM_) × Reward × Task × Response × Baseline order interaction was not significant (F(1,21) = 3.672, p > 0.05).

**Supplementary figure S2.**
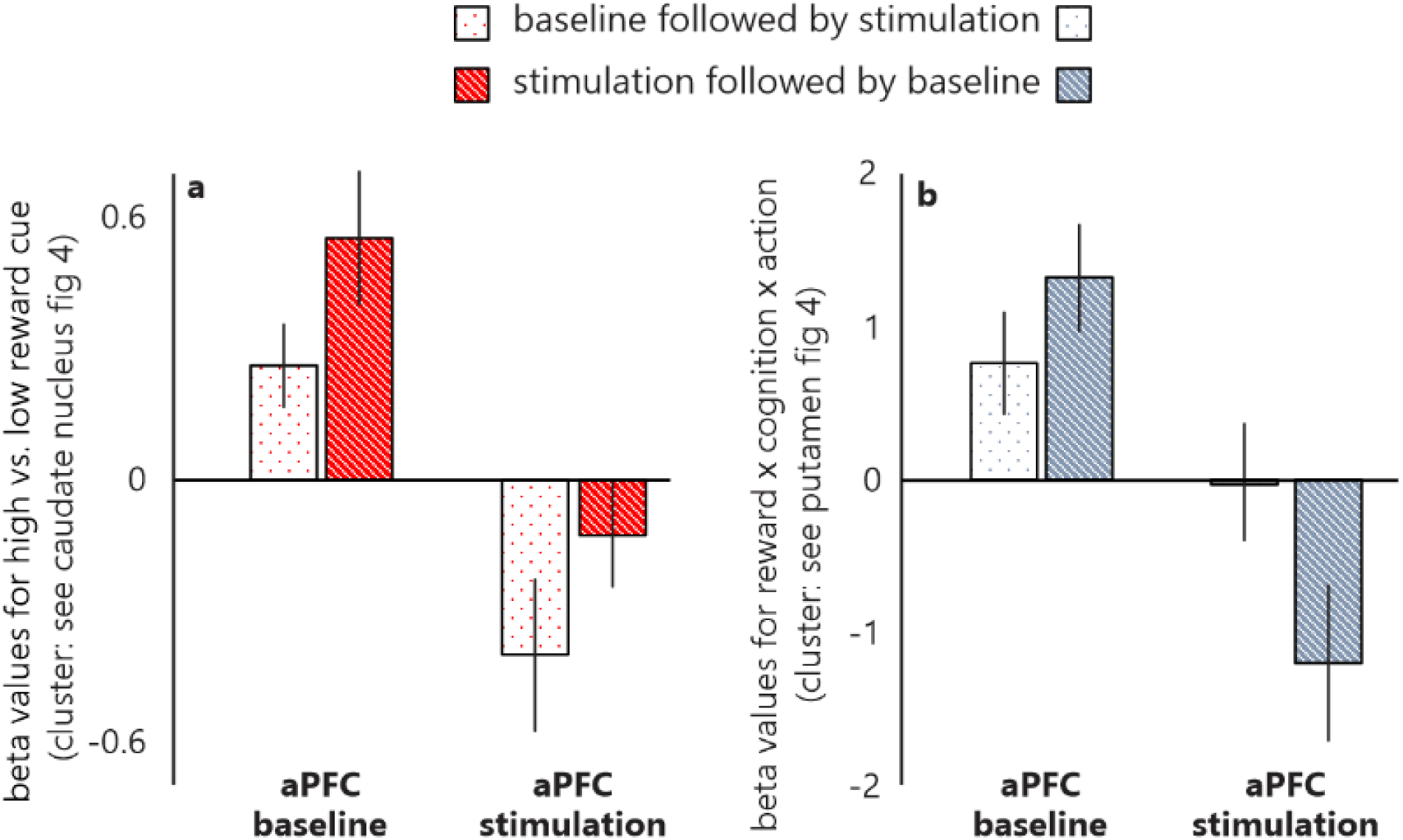
Effects of order (baseline vs. stimulation fMRI run) Plots of beta-values extracted from **a)** the right caudate nucleus cluster (see figure 4a) for the contrast high vs. low Reward and for **b)** the left putamen cluster (see figure 4b) for the contrast high vs. low Reward × Task switching (switch vs. repeat) × Response switching (switch vs. repeat). Results are shown separately for participants who performed the baseline fMRI run prior to the stimulation fMRI run (light bars) and those who performed the stimulation fMRI run before the baseline fMRI run (dark bars).

Moreover, the observed (not significant) pattern was opposite to what we would expect if the cTBS stimulation had carried over to the baseline fMRI run. If the 90 minutes between cTBS stimulation and the cTBS baseline fMRI run had not been sufficient, we would expect to see the opposite pattern from the one observed here: I.e., we would expect a smaller difference between the aPFC stimulation and the subsequent aPFC baseline session in those participants who started with the stimulation run (the lighter bars in **figure S2b**).

In addition, there was no evidence that the effect of aPFC stimulation vs. aPFC baseline on the Reward, Task, Response-related signal in the putamen (figure 4b) was dependent on the session number (1^st^, 2^nd^ or 3^rd^) in which the aPFC was stimulated (Stimulation (aPFC _BASE-STIM_ × Reward × Task × Response × Site order: F(2, 21) <1).

Finally, we repeated the analyses and included data from all sites (aPFC, dlPFC and PMC), to assess whether we could find evidence that the Site × Stimulation × Reward × Task × Response interaction in the putamen (**figure S3**) was modulated by either Stimulation order or Site order. We did not find evidence for these effects (both F’s < 1).

Inspection of **figure S2b** suggests an effect in the putamen as a function of the interaction between Reward, Task switching and Response switching during the baseline fMRI runs of the aPFC and PMC session, but not during the dlPFC session. Indeed, a direct contrast between the baseline aPFC and the baseline dlPFC session (grey bars) revealed that the BOLD signal in the left putamen was lower during the dlPFC baseline session (Site (aPFC _BASE_ vs. dlPFC _BASE_) × Reward × Task × Response: P_FWE-SVC_ = 0.019), but that there was no evidence of a difference between the baseline runs during the aPFC and PMC sessions (Site (aPFC _BASE_ vs. PMC _BASE_) × Reward × Task × Response: P_FWE-SVC_ > 0.05). To assess directly in the left putamen whether the effect in **figure S2** can be explained by a difference in BOLD response during the baseline sessions, we performed a direct test of the Reward × Task × Response × Stimulation interaction between the aPFC and PMC session (i.e. sessions with comparable baselines). This revealed a significant cluster in the left anatomical putamen (aPFC _BASE-STIM_ > PMC _BASE-STIM_ × Reward × Task × Response: P_FWE-SVC_ = 0.026, k = 26, t = 3.63, z = 3.55, x, y, z peak = −26, −8, 12). This analysis confirms that the significant 4-way interaction, reported in the main text, i.e. aPFC _BASE − STIM_ > dlPFC _BASE − STIM_ = PMC _BASE − STIM_) × Reward × Task × Response, was not driven by the dlPFC session.

**Supplementary figure S3.**
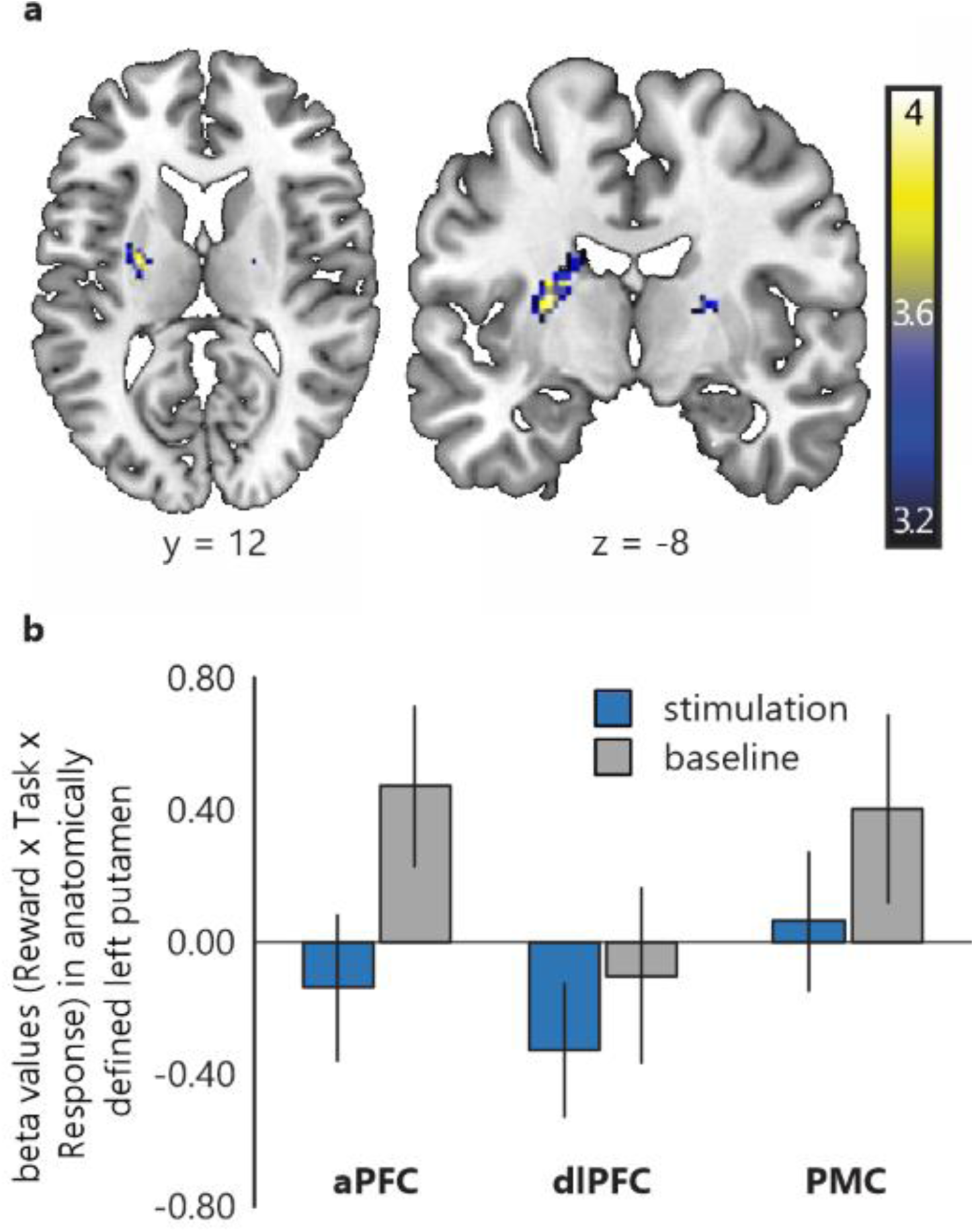
Effect of aPFC vs. dlPFC and PMC stimulation (vs. baseline) for the interaction between Reward, Task switching and Response switching. **a)** Brain maps for the interaction effect of cTBS (stimulation vs. baseline) × Site (aPFC, dlPFC, PMC) × Reward × Task switching × Response switching (shown at P_UNC_ < 0.001, note that the cluster in the left putamen is significant at P_FWE-SVC_ = 0.026). Color scales reflect t-values. **b)** Plots of the beta-values extracted from the anatomically defined left putamen for the same contrast as in figure S3a.

### Differences in the baseline run do not account for the effect of aPFC (vs. dlPFC and PMC) stimulation in the putamen

When inspecting the two sessions with equal baselines in **figure S3b**, it can be seen that the effect of aPFC _BASE-STIM_ vs. the PMC _BASE-STIM_ is driven by a combination of a numerical increase in the Reward × Task × Response –related BOLD response during the aPFC baseline fMRI run and a numerical decrease in the Reward × Task × Response –related BOLD response during the aPFC stimulation fMRI run. This was confirmed by a post-hoc test: direct comparison of the aPFC stimulation vs. PMC stimulation runs or the aPFC baseline vs. PMC baseline runs did not result in a significant cluster in the putamen. Combined, these effects resulted in a significant effect. In the absence of a baseline session, we would not have been able to detect this effect.

Thus, the inclusion of the baseline fMRI runs in our design has enabled us to take into account variation in task-related BOLD response within the same individuals across different days. This enabled us to increase the sensitivity of our measurements, quantifying inter-session variance that would otherwise have been undetectable.

### Behavioral data

#### Methods: Statistical analysis of behavioral data

The first trial of each block, trials with extremely fast responses (<100ms), and trials on which participants failed to respond were excluded from analysis. In addition, trials on which the response was incorrect were excluded from RT analyses. The RT and error rate data violated a normal distribution, which was not resolved after log and arcsine transformations, respectively (Shapiro-Wilk p < 0.05). Therefore, effects that reached significance were submitted to a non-parametric Wilcoxon signed-rank test to assure that the repeated measures GLM did not reflect false-positives. We did not observe any false-positives.

#### Results: Behavioral effects across all sessions

Across sessions, participants responded faster on high reward trials (mean ± SE: 368.07 ± 5.69ms) compared with low reward trials (mean ± SE: 375.87 ± 6.33ms) (Reward: F(1,26) = 30.930, p < 0.001, η_p_^2^= 0.543**; figure S4**). There was no main effect of reward in terms of error rates (Reward: F(1,26) < 1). In terms of task-switching performance, participants made more errors on task-switch trials (mean ± SE: 17% ± 1.6%) compared with task-repeat trials (mean ± SE: 13.7 ± 1.2%) (Task switching: F(1,26) = 25.527, p < 0.001, η_p_^2^= 0.495; **figure S4**), but showed no main effect of task switching in terms of response times (Task switching: F(1,26) = 2.146, p = 1.55). Finally, participants responded more slowly (mean ± SE: 374.26 ± 6.18ms) and made more errors (mean ± SE: 17.2 ± 1.5%) when the same response had to be repeated compared with trials on which the response switched (mean ± SE: 369.68 ± 6.14ms and 13.5 ± 1.3% respectively) (Response switching in terms of response times: F(1,26) = 8.454, p = 0.007, η_p_^2^ = 0.245), and error rates F(1,26) = 21.333, p < 0.001, η_p_^2^ = 0.451; **figure S4**).

Across sessions, participants exhibited a significant effect of reward on task switching in terms of response times (Reward × Task: F(1,26) = 40.691, p < 0.001, η_p_^2^= 0.61; **figure S4**), but not in terms of error rates (Reward × Task: F(1,26) < 1). Breaking down this effect in the response times revealed that participants exhibited a task-switch benefit (i.e. task repeat (mean ± SE:377.73 ± 6.23ms) – task switch (mean ± SE: 374.00 ± 6.51ms)) on low reward trials (F(1,26) = 7.805, p = 0.010, η_p_ ^2^= 0.231) and a switch cost (i.e. task switch (mean ± SE: 371.57 ± 6.28ms) – task repeat (mean ± SE: 364.58 ± 5.71ms) performance) on high reward trials (F(1,26) = 23.305, p< 0.001, η_p_ ^2^= 0.473). In addition, in terms of errors rates, we observed a larger task-switch cost on response repetition trials (task switch: mean ± SE: 19.7% ± 1.8%; task repeat: mean ± SE: 14.6% ± 1.4%; main effect of task: F(1,26) = 25.910, p < 0.001, η_p_ ^2^= 0.499) than on response switch trials (task switch: mean ± SE: 14.3% ± 1.5%; task repeat: mean ± SE: 12.8% ± 1.2%; main effect of task: F(1,26) = 4.266, p = 0.049, η_p_ ^2^= 0.141; **figure S4**). This was evidenced by a significant Task × Response interaction (error rates: F(1,26) = 9.489, p = 0.005, η_p_ ^2^= 0.267; response times: F(1,26) <1). There was no Reward × Task × Response interaction (F(1,26) < 1) for response times or error rates).

### Discussion of behavioral data

A number of studies have reported that task-switching and response switching are not independent^1,2^. This is substantiated by our behavioral data. The task-switch cost observed was larger on response-repeat trials, in agreement with previous reports^1,2^. In addition, we observed a behavioral response switch benefit: participants were faster and more accurate when responding on a response switch compared with a response repeat trial **(figure S4)**. When inspecting **figure S4**, it becomes clear that there was a considerable behavioral cost associated with trials on which the task switched, but the response remained the same. It appears that this increased error rate on response repeat/task switch trials is driving both the interaction and the main effect.

**Supplementary figure S4.**
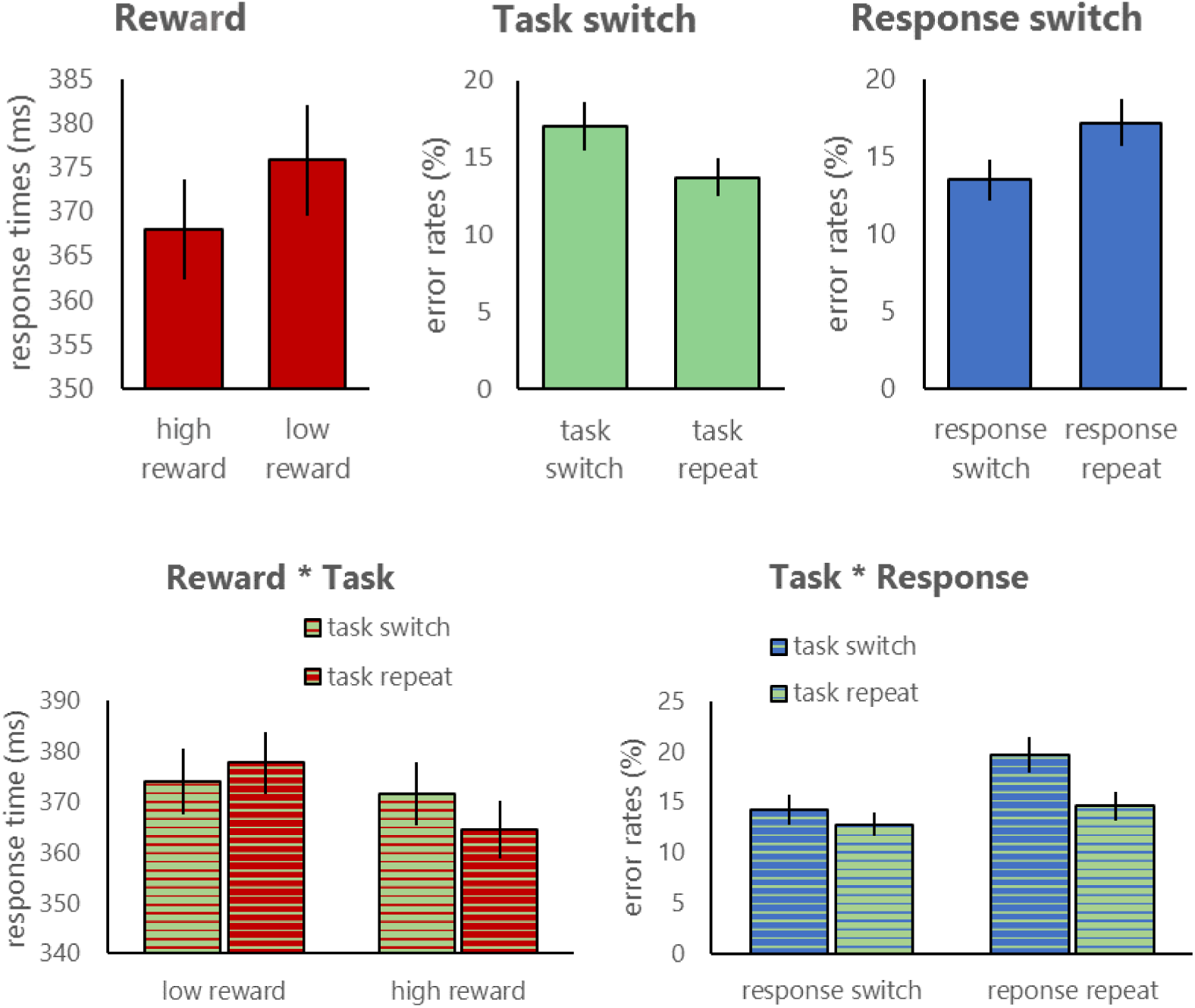
Behavioral data across all sessions. Data are shown for **top:** the main effect of reward (response times), task switching (error rates) and response switching (error rates), and **bottom:** for the interaction between reward and task switching (Reward × Task; response times) and for the interaction between task switching and response switching (Task × Response; error rates). All effects are shown across all 6 runs, i.e. irrespective of TMS.

## References

1. Alexander, G. E., DeLong, M. R. & Strick, P. L. Parallel organization of functionally segregated circuits linking basal ganglia and cortex. Annu. Rev. Neurosci. 9, 357–381 (1986).

2. Alexander, G. E. & Crutcher, M. D. Functional architecture of basal ganglia circuits: neural substrates of parallel processing. TINS 13, 266–271 (1990).

3. Draganski, B. et al. Evidence for Segregated and Integrative Connectivity Patterns in the Human Basal Ganglia. J. Neurosci. 28, 7143–7152 (2008).

4. Di Martino, A. et al. Functional Connectivity of Human Striatum: A Resting State fMRI Study. Cereb. Cortex 18, 2735–2747 (2008).

5. Cromwell, H. C. & Schultz, W. Effects of Expectations for Different Reward Magnitudes on Neuronal Activity in Primate Striatum. J. Neurophysiol. 89, 2823–2838 (2003).

6. Stelzel, C., Basten, U., Montag, C., Reuter, M. & Fiebach, C. J. Frontostriatal Involvement in Task Switching Depends on Genetic Differences in D2 Receptor Density. J. Neurosci. 30, 4205–14212 (2010).

7. Lehericy, S. et al. Motor control in basal ganglia circuits using fMRI and brain atlas approaches. Cereb. Cortex16, 149–161 (2006).

8. Haber, S. N., Fudge, J. L. & McFarland, N. R. Striatonigrostriatal pathways in primates form an ascending spiral from the shell to the dorsolateral striatum. J. Neurosci. 20, 2369–2382 (2000).

9. Haber, S. N. The primate basal ganglia: parallel and integrative networks. J. Chem. Neuroanat. 26, 317–330 (2003).

10. Haber, S. N., Kim, K.-S., Mailly, P. & Calzavara, R. Reward-Related Cortical Inputs Define a Large Striatal Region in Primates That Interface with Associative Cortical Connections, Providing a Substrate for Incentive-Based Learning. J. Neurosci. 26, 8368–8376 (2006).

11. Marquand, A. F., Haak, K. V. & Beckmann, C. F. Functional corticostriatal connection topographies predict goal-directed behaviour in humans. 1, 0146 (2017).

12. Jarbo, K. & Verstynen, T. D. Converging Structural and Functional Connectivity of Orbitofrontal, Dorsolateral Prefrontal, and Posterior Parietal Cortex in the Human Striatum. J. Neurosci. 35, 3865–3878 (2015).

13. Strafella, A. P., Paus, T., Barrett, J. & Dagher, A. rRepetitive transcranial magnetic stimulation of the human prefrontal cortex induces dopamine release in the caudate nucleus. J Neurosci21, 1–4 (2001).

14. Strafella, A. P., Paus, T., Fraraccio, M. & Dagher, A. Striatal dopamine release induced by repetitive transcranial magnetic stimulation of the human motor cortex. Brain 126, 2609–2615 (2003).

15. Ko, J. H. et al.Theta burst stimulation-induced inhibition of dorsolateral prefrontal cortex reveals hemispheric asymmetry in striatal dopamine release during a set-shifting task - a TMS-[ ^11^ C]raclopride PET study. Eur. J. Neurosci. 28, 2147–2155 (2008).

16. Hanlon, C. A. et al.Probing the Frontostriatal Loops Involved in Executive and Limbic Processing via Interleaved TMS and Functional MRI at Two Prefrontal Locations: A Pilot Study. PLoS ONE 8, (2013).

17. van Schouwenburg, M. R., O’Shea, J., Mars, R. B., Rushworth, M. F. S. & Cools, R. Controlling Human Striatal Cognitive Function via the Frontal Cortex. J. Neurosci. 32, 5631–5637 (2012).

18. Zandbelt, B. B., Bloemendaal, M., Hoogendam, J. M., Kahn, R. S. & Vink, M. Transcranial magnetic stimulation and functional MRI reveal cortical and subcortical interactions during stop-signal response inhibition. J. Cogn. Neurosci. 25, 157–174 (2013).

19. Huang, Y.-Z., Edwards, M. J., Rounis, E., Bhatia, K. P. & Rothwell, J. C. Theta Burst Stimulation of the Human Motor Cortex. Neuron45, 201–206 (2005).

20. Volman, I., Roelofs, K., Koch, S., Verhagen, L. & Toni, I. Anterior Prefrontal Cortex Inhibition Impairs Control over Social Emotional Actions. Curr. Biol. 21, 766–1770 (2011).

21. Wischnewski, M. & Schutter, D. J. L. G. Efficacy and Time Course of Theta Burst Stimulation in Healthy Humans. Brain Stimulat. 8, 685–692 (2015).

22. Aarts, E. et al. Striatal Dopamine Mediates the Interface between Motivational and Cognitive Control in Humans: Evidence from Genetic Imaging. Neuropsychopharmacology 35, 1943–1951 (2010).

23. van Holstein, M. et al.Human cognitive flexibility depends on dopamine D2 receptor signaling. Psychopharmacology (Berl.)218, 567–578, (2011).

24. Aarts, E. et al.Greater striatal responses to medication in Parkinson?s disease are associated with better task-switching but worse reward performance. Neuropsychologia62, 390–397 (2014).

25. Aarts, E. et al.Reward modulation of cognitive function in adult attention-deficit/hyperactivity disorder: a pilot study on the role of striatal dopamine. Behav. Pharmacol. 26, 227–240 (2015).

26. Piray, P., den Ouden, H. E. M., van der Schaaf, M. E., Toni, I. & Cools, R. Dopaminergic Modulation of the Functional Ventrodorsal Architecture of the Human Striatum. Cereb. Cortex(2015). doi:10.1093/cercor/bhv243

27. Schutter, D. L. G. & van Honk, J. A standardized motor threshold estimation procedure for transcranial magnetic stimulation research. J. ECT 22, 176–178 (2006).

28. Poser, B. A., Versluis, M.J., Hoogduin, J. M.& Norris, D. G. BOLD contrast sensitivity enhancement and artifact reduction with multiecho EPI: parallel-acquired inhomogeneity-desensitized fMRI. Magn. Reson. Med. 55, 1227–1235 (2006).

29. Haber, S. N. & Knutson, B. The reward circuit: linking primate anatomy and human imaging. Neuropsychopharmacol. Off. Publ. Am. Coll. Neuropsychopharmacol. 35, 4–26 (2010).

30. Taber, M. T., Das, S. & Fibiger, H. C. Cortical regulation of subcortical dopamine release: mediation via the ventral tegmental area. J. Neurochem. 65, 1407–1410 (1995).

31. Ferreira, J. G. P., Del-Fava, F., Hasue, R. H. & Shammah-Lagnado, S. J. Organization of ventral tegmental area projections to the ventral tegmental area-nigral complex in the rat. Neuroscience153, 196–213 (2008).

32. Fitzpatrick, S. M. & Rothman, D. L. Meeting report: transcranial magnetic stimulation and studies of human cognition. J. Cogn. XsNeurosci. 12, 704–709 (2000).

33. Pell, G. S., Roth, Y. & Zangen, A. Modulation of cortical excitability induced by repetitive transcranial magnetic stimulation: influence of timing and geometrical parameters and underlying mechanisms. Prog. Neurobiol. 93, 59–98 (2011).

34. Derrfuss, J., Brass, M., Neumann, J. & von Cramon, D. Y. Involvement of the inferior frontal junction in cognitive control: meta-analyses of switching and Stroop studies. Hum Brain Mapp25, 22–34 (2005).

35. Stelzel, C., Basten, U. & Fiebach, C. J. Functional connectivity separates switching operations in the posterior lateral frontal cortex. J. Cogn. Neurosci. 23, 3529–3539 (2011).

36. Stelzel, C., Fiebach, C. J., Cools, R., Tafazoli, S. & D’Esposito, M. Dissociable fronto-striatal effects of dopamine D2 receptor stimulation on cognitive versus motor flexibility. Cortex J. Devoted Study Nerv. Syst. Behav. 49, 2799–2811 (2013).

37. Tupak, S. V. et al.Inhibitory transcranial magnetic theta burst stimulation attenuates prefrontal cortex oxygenation. Hum. Brain Mapp.34, 150–157 (2013).

38. Volkow, N. D. & Fowler, J. S. Volkow Fowler 2000 Addiction a disease of compulsion and drive.pdf.Cereb. Cortex10, 318–325 (2000).

39. Shepherd, G. M. G. Corticostriatal connectivity and its role in disease. Nat. Rev. Neurosci. 14, 278–291 (2013).

40. Malone, D. A. et al.Deep brain stimulation of the ventral capsule/ventral striatum for treatment-resistant depression. Biol. Psychiatry65, 267–275 (2009).

41. Denys, D. et al.Deep brain stimulation of the nucleus accumbens for treatment-refractory obsessive-compulsive disorder. Arch Gen Psychiatry67, 1061–8 (2010).

42. Weaver, F. M. et al.Bilateral deep brain stimulation vs best medical therapy for patients with advanced Parkinson disease: a randomized controlled trial. JAMA301, 63–73 (2009).

43. Dlabac-de Lange, J. J., Knegtering, R. & Aleman, A. Repetitive transcranial magnetic stimulation for negative symptoms of schizophrenia: review and meta-analysis. J. Clin. Psychiatry71, 411–418 (2010).

## Supplementary references

1. Shook, S. K., Franz, E. A.,Higginson, C. I., Wheelock, V. L. & Sigvardt, K. A. Dopamine dependency of cognitive switching and response repetition effects in Parkinson’s patients. Neuropsychologia 43, 1990–1999(2005)

2. Stelzel, C, Basten, U & Fiebach, C.J Functional connectivity separates switching operations in the posterior lateral frontal cortex. J. Cogn. Neurosci 23, 3529–3539(2011).

